# Electrical Control of the Transduction Channels’ Gating Force in Mechanosensory Hair Cells

**DOI:** 10.1101/2024.12.13.628311

**Authors:** Achille Joliot, Laure Stickel, Pascal Martin

## Abstract

The inner ear’s hair cells rely on mechanosensitive ion channels to convert vibrations of their hair bundles into electrical signals. The mechanical correlates of channel gating—the gating force and the gating swing—are fundamental determinants of hair-cell mechanosensitivity but are still poorly understood. Here we show that varying the electrical potential across the sensory hair-cell epithelium continuously modulates the gating force, by up to ±100%. Our observations also revealed an abrupt transition between states of weak and strong gating force at a threshold potential, so that strong gating forces associated to high mechanosensitivity are observed only when the calcium influx through the channels is large enough, but not too large. Gating-force changes, remarkably enough, were explained by the modulability of the gating swing, ranging from values comparable to the channel pore size to nearly tenfold larger. Gating-swing control is expected to underly the hair cell’s ability to tune its mechanosensitivity to minute sound stimuli.

**Table of content:** The inner ear’s hair cells rely on mechanosensitive ion channels to convert vibrations of their hair-bundle into electrical signals. We show that varying the electrical potential (*U*) across the sensory epithelium modulates a key determinant of mechanosensitivity—the gating force (*F*_*G*_)—by modulating the gating swing (*d*), ranging from the size of the channels’ pore to nearly tenfold larger.

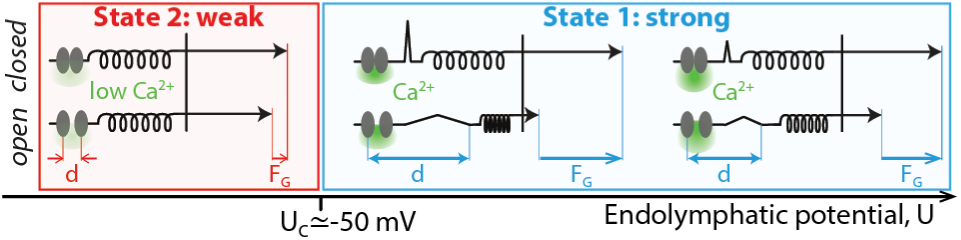

## 1. Introduction

Mechanoreception by the inner ear’s sensory hair cells begins with the deflection of their hair bundle—a cohesive tuft of cylindrical stereocilia arranged in a staircase pattern.^[1–3]^ A deflection toward the taller row of stereocilia increases mechanical tension in oblique tip links, which connect the tip of each stereocilium to the flank of a taller neighbor. The increased tip-link tension opens mechanosensitive ion channels and evokes an influx of cations into the hair cell, primarily K^+^ ions but also Ca^2+^ ions.

Our understanding of hair-cell mechanosensitivity is informed by the gating-spring model of mechanoelectrical transduction.^[4,5]^ Within this framework, the transduction channels are gated directly by force, which is transmitted to the channels via elastic elements—the gating springs—that operate in series with the tip links. The product of the gating-spring stiffness and the gating swing defines the gating force, a fundamental determinant of hair-cell mechanosensitivity: its magnitude sets the maximal slope of the sigmoidal relation between the transduction current and the deflection of the hair bundle (Figure S1A,B, Supporting Information). The greater the gating force, the larger the transduction current evoked by a small stimulus, making the hair cell more sensitive to a given deflection of its hair bundle.

The gating-spring model imposes a reciprocal relationship between tip-link tension and channel gating. If tensioning the tip links evokes channel opening, then channel opening must reduce tip-link tension by an amount given precisely by the gating force. Feedback between tip-link tension and channel gating produces gating compliance, the effective softening of the hair bundle upon channel gating.^[6,7]^ For large positive or negative bundle displacements, at which the transduction channels are nearly all open or closed, respectively, the force-displacement relation is linear. At intermediate displacements that elicit channel gating, however, the slope of the force-displacement relation is reduced: the hair bundle is effectively softer. Gating compliance is associated with a shift between the two linear limbs of the force-displacement relation; this shift provides an estimate of the combined gating force, denoted *F*_*G*_in the following, produced by the ensemble of transduction channels that operate in parallel within the hair bundle (Figure S1B, Supporting Information). Furthermore, upon an oscillatory movement of the hair bundle, opening and closing of the transduction channels also increase friction on the hair bundle, a phenomenon called gating friction.^[8]^ The experimental demonstration of gating compliance and gating friction at the cellular scale of a micron-sized hair bundle provides strong support for the gating-spring model. Because these phenomena reflect the mechanical correlate of channel gating, they can also be used to infer fundamental properties of channel gating at the molecular scale such as the gating force.^[6,7,9]^

Biophysical estimates of the gating force are variable, even with hair cells from the same animal species and organ. With hair cells in excised preparations of the mammalian cochlea, there is some evidence of gating compliance ^[10–14]^ but gating forces can also be so weak that the force-displacement relation remains linear.^[15–17]^ In these experiments, however, the endocochlear potential that normally boosts transduction currents in vivo is absent and the hair cells are fully immersed in a standard extracellular fluid called perilymph, which is rich in Na^+^ and with a Ca^2+^ concentration in the millimolar range. This condition disrupts the two ionic compartments that hair cells experience in vivo, with endolymph—an unusual K^+^-rich fluid with a low Ca^2+^ concentration (20-250 µM)—and perilymph bathing the cells’ apical and basal aspects, respectively. In addition, because cochlear hair bundles are weakly cohesive, gating-force estimates depend on the method of mechanical stimulation ^[17,18]^ and where displacement is measured within the bundle.^[19]^

The frog saccule has provided a robust model system to study gating forces. Hair bundles are highly cohesive, which ensures concerted gating of the transduction channels of their hair bundles.^[20,21]^ In addition, because adaptation is relatively slow, the effects of channel gating on hair-bundle mechanics can be evaluated before adaptation kicks in and obscures the interpretation of force measurements.^[9]^ Finally, the organ can be mounted in a two-chamber configuration, which allows to mimic the natural ionic conditions. ^[22,23]^ With frog saccular hair cells, gating forces can lead to a moderate (about 40%) decrease in hair-bundle stiffness ^[6]^ or be strong enough to result in negative stiffness, which can lead to spontaneous oscillations of the hair bundle and, in turn, frequency-selective amplification of periodic stimuli.^[9,23–25]^ Remarkably enough, strong gating forces and negative stiffness were reported only under ex-vivo conditions that closely mimic natural ones, with reconstituted endolymph-perilymph compartments. This and other circumstantial evidence ^[26,27]^ suggests that the magnitude of the gating force may vary with the Ca^2+^ concentration in the fluid bathing the hair bundle.

In this work, we clarify the apparent variability of the gating force by providing a comprehensive and systematic investigation of the mechanical correlates of channel gating in hair-bundle mechanics under a range of conditions. We report that the electrical potential across the sensory epithelium that houses the hair cells can serve as a control parameter of the gating force. Our results indicate that the gating force varies primarily because the gating swing varies. Our work further reveals a reversible, but hysteretic, transition between a state of strong gating force, for which the hair bundles oscillate spontaneously, and a state of weak gating force, for which hair-bundle mechanics is linear and the hair bundles are quiescent. We finally provide evidence that these effects are mediated, at least in part, by electrical control of the influx of Ca^2+^ through the transduction channels.

## 2. Results

We took advantage of an excised preparation of the frog saccule that mimics *ex vivo* the two-compartment ionic conditions that the sensory epithelium experiences *in vivo*. Under these conditions, the hair cells routinely displayed spontaneous oscillations of their hair bundles, as recognized before.^[26,28,29]^ We aimed here at evaluating the effects of a transepithelial potential on hair-bundle mechanics. To this end, we used a pair of electrodes and a current generator to impose a transepithelial current, *I*, within a range of ±10 µA (**Figure** 1A). Upon application of a current step, the transepithelial potential changed with a time course that was well described by an exponential relation with a characteristic timescale of 230 ± 50 μs (mean ± SD, *n* =13; Figure 1B). The transepithelial potential thus reached a steady-state value, noted *U* in the following, in about 600 µs. Over an ensemble of 13 saccules, we found a transepithelial resistance of 10 ± 2 kΩ (mean ± SD). Thus, a positive current of +1 µA evoked a positive potential in the endolymphatic compartment of about +10 mV with respect to the perilymphatic compartment; accordingly, the transepithelial potential thus varied within a range of ±100 mV. We emphasize for the following that a positive endolymphatic potential is expected to increase the influx of K^+^ and Ca^2+^ ions through the transduction channels, whereas a negative potential does the opposite. All numerical results are given below as the average and standard deviations (SD) over an ensemble of *n* independent measurements.

**Figure 1:**
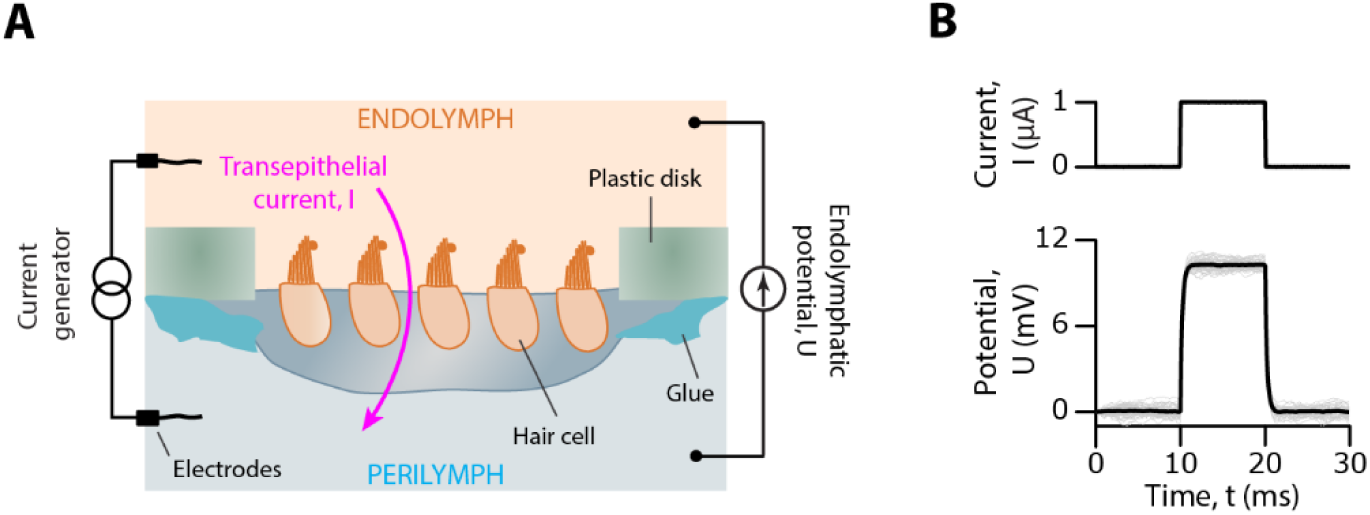
Application of an endolymphatic potential in a two-compartment ex-vivo preparation of the frog saccule. **(A)** Schematic of the setup. The tissue was glued over a 1-mm hole in a plastic disk. Its apical and basal aspects were bathed in artificial endolymph and perilymph, respectively. A transepithelial current, *I*, was produced by a pair of silver chloride electrodes connected to a current generator, resulting in an endolymphatic potential, *U*, with respect to the potential in perilymph. The sign convention is such that positive currents lead to a positive value of the endolymphatic potential. **(B)** In this representative example, a +1-µA step of transepithelial current (top) evoked an endolymphatic potential *U* ≃ +10 mV at steady state (bottom, grey), corresponding to a transepithelial resistance of 10 kΩ. An exponential fit (black line), *U*(*t*) = *A* exp(−t/τ), to the response yielded the amplitude *A* = 10.2 mV and the characteristic time *τ* = 210 μs.

### 2.1 The potential modulates strongly the gating force but weakly the hair-bundle stiffness

To determine the effects of an endolymphatic potential on the mechanical properties of a hair bundle, we measured the bundle’s force-displacement relation in response to a series of step displacements applied to the base of a stimulus fiber (**Figure 2**). Under control conditions, for which no transepithelial current was applied (*I* = 0), the force-displacement relation showed a prominent region of reduced slope, or gating compliance, which is expected for hair bundles that oscillate spontaneously.^[9,24]^ A fit of the relation to Equation 1 (Experimental Section) yielded a set-point deflection X_0_ = −2 ± 7 nm, a gating force *F*_*G*_ = 17 ± 4 pN and a linear stiffness *K* = 0.85 ± 0.2 pN ⋅ nm^−1^ (*n* = 29 in 16 frogs). We recall that the set-point deflection can be interpreted as the deflection that leads to a channels’ open probability of ½.^[24]^ For each hair bundle, the procedure was repeated in the presence of a static transepithelial current (*I* ≠ 0).

**Figure 2:**
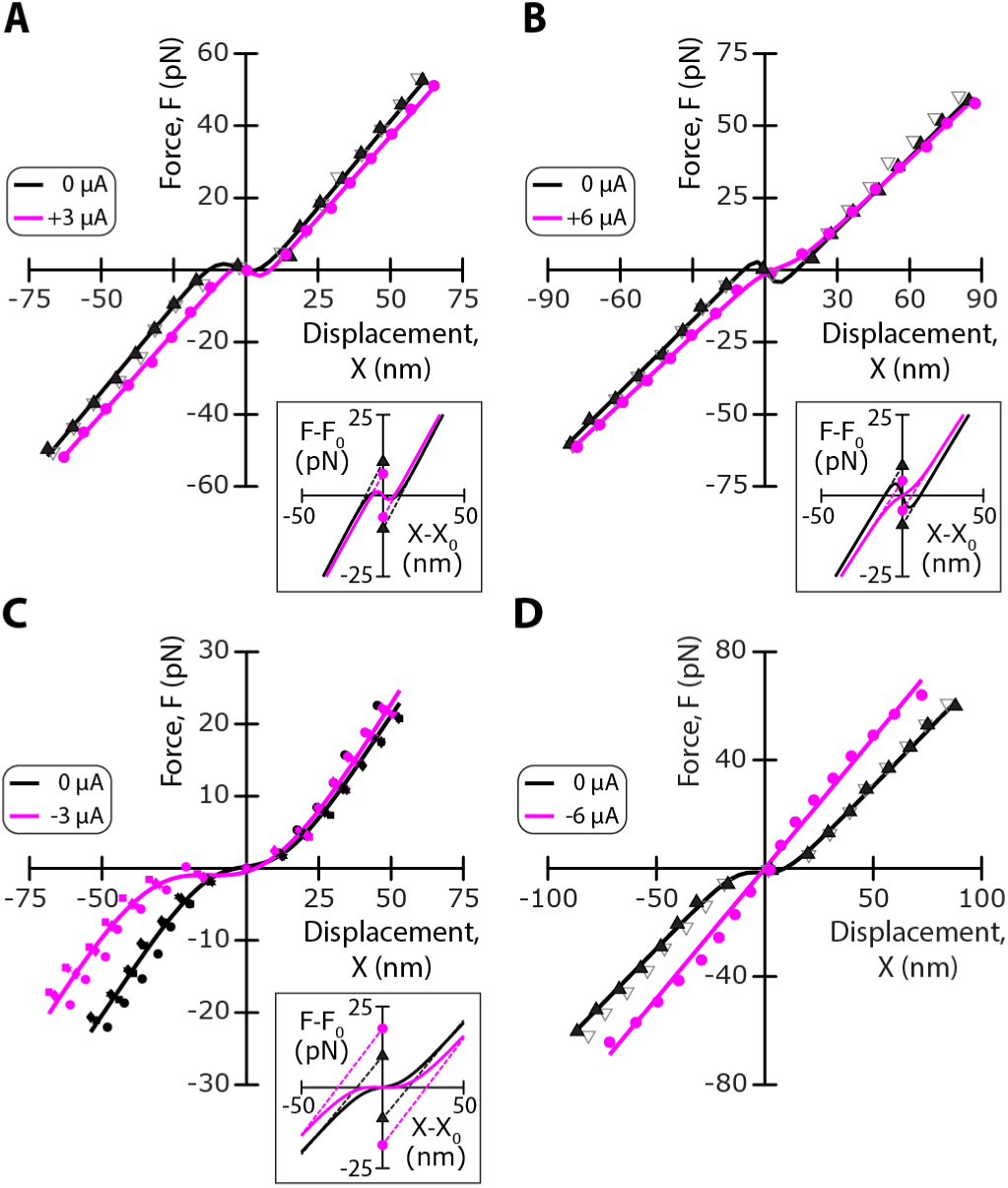
Representative examples of mechanical changes evoked by a transepithelial current. **(A-D)** Force-displacement relations, *F*(X), for four different hair bundles under control conditions (*I* = 0 μA; black) and in the presence of a transepithelial current (magenta) of amplitude *I* = +3 μA (A), *I* = +6 μA (B), *I* = −3 μA (C) and *I* = −6 μA (D). Each relation was fitted by Equation 1 (solid lines; parameter values in Table S1). Insets: the fitted curves are replotted with respect to the origin (X_0_, *F*(X_0_)), where X_0_ is the set-point deflection. Under each condition, the hair bundle’s gating force, *F*_*G*_, was given by the vertical distance between the two symbols in the inset (*I* = 0: triangles; *I* ≠ 0: disks). In (A), (B) and (D), the open black symbols correspond to a measurement after restoring control conditions to assess reversibility; in (C), the experiment was repeated 4 times.

We start by presenting the effects of a transepithelial current on a force-displacement relation in four representative examples (Figure 2); the effects on the corresponding spontaneous hair-bundle oscillations are shown in Figure S2A-D (Supplementary Information). Applying a positive current, here *I* = +3 μA, corresponding to a positive endolymphatic potential *U* ≃ +30 mV, evoked an increase of the set-point deflection: the transduction channels closed (Figure 2A). In addition, we observed a small (−5%) decrease of the linear bundle’s stiffness and, remarkably, a large decrease (−38%) of the gating force. In another cell, a larger current *I* = +6 μA resulted in a similar change in stiffness (−6%) but evoked a larger decrease of the gating force (−50%; Figure 2B). Conversely, applying a negative current of moderate magnitude, here *I* = −3 µA, corresponding to a negative endolymphatic potential U ≃ −30 mV, elicited a decrease of the set-point deflection and thus opening of the transduction channels, a small increase (+10%) of the linear stiffness, and a large increase (+89%) of the gating force (Figure 2C). Furthermore, applying a large transepithelial current, here I = −6 µA, led to an unexpected finding: the force-displacement relation became linear (Figure 2D). This is not because the integrity of the hair cell was impaired by application of a large transepithelial current, for all the effects were fully reversible.

Performing an ensemble of *n* = 41 pairs of measurements in 29 hair bundles from 16 frogs confirmed the effects illustrated by the four representative examples. We found a strong positive correlation between the change, *Δ*X_0_ = X_0_(*I*) − X_0_(*I* = 0), of the set-point deflection and the applied transepithelial current (Pearson correlation r = 0.87; p-value: 4 · 10^−11^; *n* = 34; **Figure 3A**). This correlation came as no surprise, for the endolymphatic potential resulting from the transepithelial current should influence the magnitude of the Ca^2+^ influx through the transduction channels, and calcium is known to promote channel closure.^[4,24,27,30,31]^ The change in linear stiffness, *ΔK* = *K*(*I*) − *K*(*I* = 0), was negatively correlated to the transepithelial current, but the correlation was weaker (Pearson correlation r = −0.57 ; p-value: 10^−4^; *n* = 41) (Figure 3B). Instead, for currents *I* > −4 µA, the change, *ΔF*_*G*_ = *F*_*G*_(*I*) − *F*_*G*_(*I* = 0), of the hair-bundle’s gating force displayed a strong negative correlation with the current (Pearson correlation r = −0.86 ; p-value: 8 · 10^−11^; *n* = 34; Figure 3C). The relative variation of the gating force was correspondingly much larger than that of the linear stiffness (Figure 3D). A positive transepithelial current evoked a reduction of the gating force by as much as −80%, whereas the stiffness decreased at most by −10% over the same range of transepithelial currents. Conversely, the gating force could nearly double in the presence of a negative transepithelial current, corresponding to a relative increase of +100%, whereas stiffness did not increase beyond +25%. These correlations between mechanical properties of the hair bundle and the transepithelial current were associated to systematic changes of the frequency, amplitude and waveform of spontaneous hair-bundle oscillations (Figure S2E-G, Supplementary Information).

**Figure 3:**
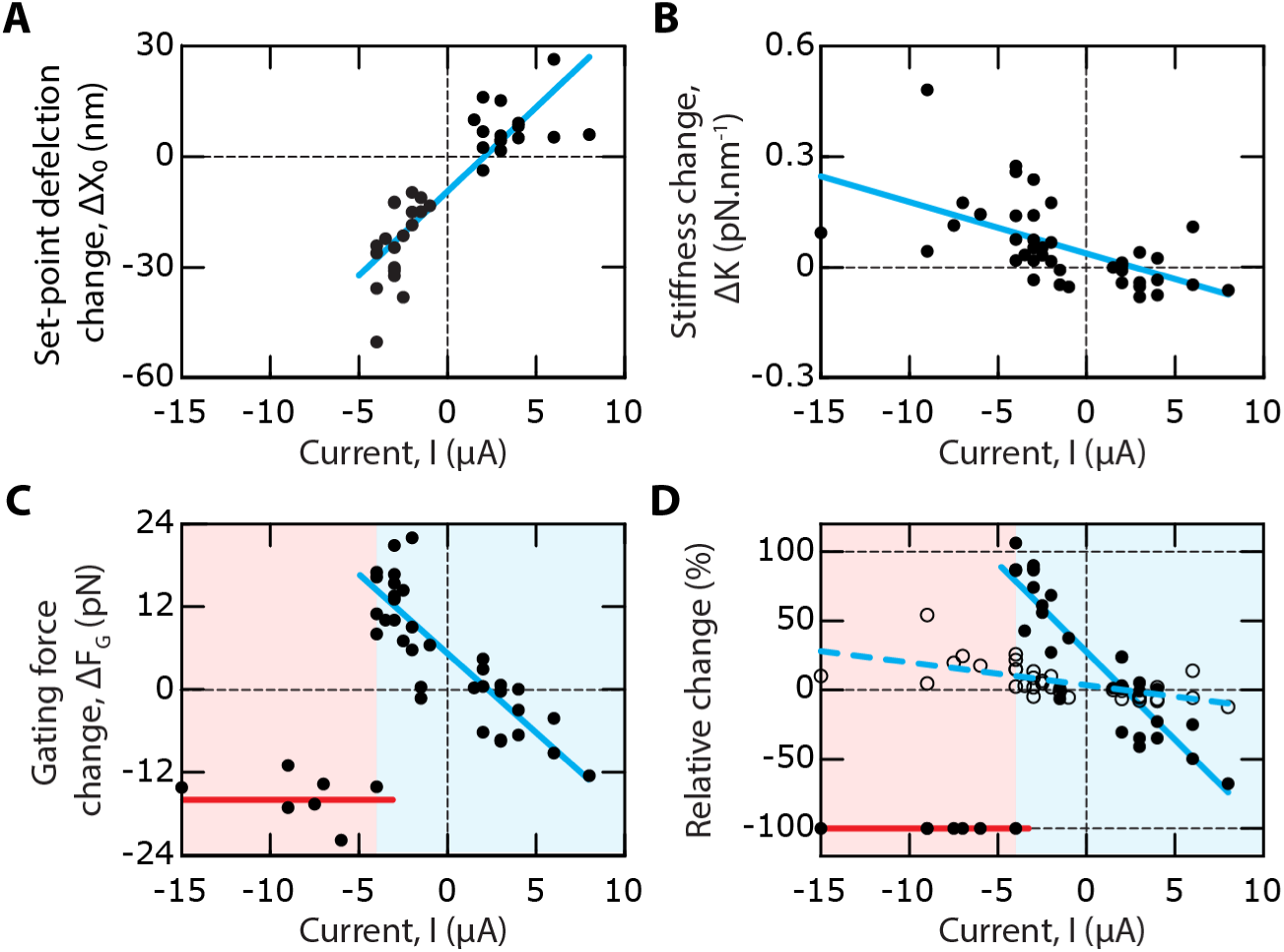
Changes in set-point deflection, stiffness and gating force upon application of a transepithelial current. **(A)** The change, *Δ*X_0_, of the set-point deflection showed a strong positive correlation to the transepithelial current, *I* (Pearson correlation r = 0.87; p-value: 4 · 10^−11^; *n* = 34). Linear regression (cyan line): ΔX_0_ [nm] = 4.6 *I* [µA] − 9.6 (*R*^2^ = 0.75). The set-point deflection could only be measured for I ≥ −5 μA. **(B)** The change, *ΔK*, of the linear stiffness was negatively correlated to the current (Pearson correlation r = −0.57 ; p-value: 10^−4^; *n* = 41). Linear regression (cyan line): Δ*K* [pN . nm^−1^] = −0.01 *I* [µA] + 0.04 (*R*^2^ = 0.33) **(C)** For I ≥ −5 μA (cyan area) the change, *ΔF*_*G*_, of the bundle’s gating force showed a strong negative correlation to the current (Pearson correlation r = −0.86 ; p-value: 8 · 10^−11^; *n* = 34). Linear regression (cyan line): Δ*F*_*G*_ [pN] = −2.3 *I* [µA] + 5 (*R*^2^ = 0.74). For I < −5 μA (red area), all the force displacement relations were linear: the gating force had dropped to undetectable levels. Accordingly, the mean value of the change Δ*F*_*G*_ = −16 ± 3 pN (mean ± SD, *n* = 7; red line) had an absolute magnitude near that of the gating force, *F*_*G*_(*I* = 0) = 17 ± 4 pN (mean ± SD, *n* = 29) under control conditions. **(D)** Relative change of the hair-bundle’s gating force, Δ*F*_*G*_/*F*_*G*_(0) (solid disks) and linear stiffness, Δ*K*/*K*(0) (open disks) as a function of the transepithelial current, *I*. Linear regressions: Δ*F*_*G*_/*F*_*G*_(0) [%] = −13 · *I* [µA] + 28 (*R*^2^ = 0.8; cyan solid line) and Δ*K*/*K*(0) [%] = −1.6 *I* [µA] + 3.0 (*R*^2^ = 0.32; cyan dashed line). For *I* ≤ −5 µ*A*, Δ*F*_*G*_/*F*_*G*_(0) = −100 % (red line).

For all the seven hair bundles for which the transepithelial current was more negative than −5 µA, corresponding to an endolymphatic potential *U* ≤ −50 mV, the gating force did not follow the trend observed at more positive potentials. Instead of increasing further or saturating at more negative potentials, the gating force dropped to values so low that the force-displacement relation was linear, thus with no detectable signature of gating compliance (Figures 2D and 3C,D). At these endolymphatic potentials, no spontaneous oscillation was observed; the hair bundles were quiescent (Figure S2D, Supplementary Information). Our study thus revealed the existence of a critical value of the transepithelial current, *I* = *I*_*C*_ ≃ −5 µ*A*, and of the endolymphatic potential, *U* = *U*_*C*_ ≃ −50 mV, that delimited two functional states of the hair bundle: a state of strong gating force for which gating compliance was large enough to yield spontaneous oscillations of the hair bundle and a state of weak gating force for which hair-bundle mechanics was linear and the hair bundle was quiescent.

### 2.2 The gating force varies because the gating swing varies

Within the framework of the gating-spring model (Figure S1A, , Supplementary Information), the gating force for the whole hair bundle, *F*_*G*_ = *K*_*GS*_*D*, is the product of the combined stiffness of the gating springs, *K*_*GS*_, and of the gating swing, *D*, of a transduction channel. We note that this interpretation of the gating force rests on the tight cohesiveness of hair bundles in the frog saccule,^[20,21]^ so that force directed to the transduction channels upon deflection of the bundle’s tip is distributed evenly across tip links.^[32]^ Remarkably enough, the relative effect of an endolymphatic potential on the gating force was much larger than that on the hair-bundle stiffness and could even be of opposite sign (Figure 3D). The contrast in behavior was particularly striking at the transition between regimes of strong and weak gating force ( *I* ≤ *I*_*C*_). Although the gating force dropped to undetectable levels, the hair-bundle stiffness instead showed a mild increased with respect to control conditions. Because the gating springs are expected to contribute a significant fraction of the total hair-bundle stiffness,^[33]^ *K*, we reasoned that the change in gating force resulted mainly from a change in the magnitude of the gating swing. We could confirm this inference by measuring the hair-bundle stiffness, *K*_*SP*_ = 0.17 ± 0.03 pN ⋅ nm^−1^ (*n* = 14) ≃ 0.2 *K*, after disruption of the tip links. By subtracting the average value of *K*_*SP*_ to individual measurements of *K*, we could then estimate, for each hair bundle, the gating-spring stiffness *K*_*GS*_ = *K* − *K*_*SP*_, and in turn the gating swing *D*. After projection along the oblique axis of the tip links using the projection factor *γ* = 0.14,^[32]^ we found a gating swing *d* = *γD* = 3.7 ± 1.4 nm (*n* = 29) under control conditions (**Figure 4A**). Replicating the behavior of the gating force, the gating-swing change displayed a strong negative correlation with the transepithelial current for currents *I* ≥ *I*_*C*_ (Pearson correlation *r* = −0.86; p-value = 7 ⋅ 10^−11^; *n* = 34) and dropped to undetectable levels for *I* ≤ *I*_*C*_ (Figure 4B). For *I* ≥ *I*_*C*_, the gating swing *d* could reach a value of up to about 8 nm in the presence of a negative endolymphatic potential or, conversely, be as small as 2 nm upon application of a potential of opposite polarity.

**Figure 4:**
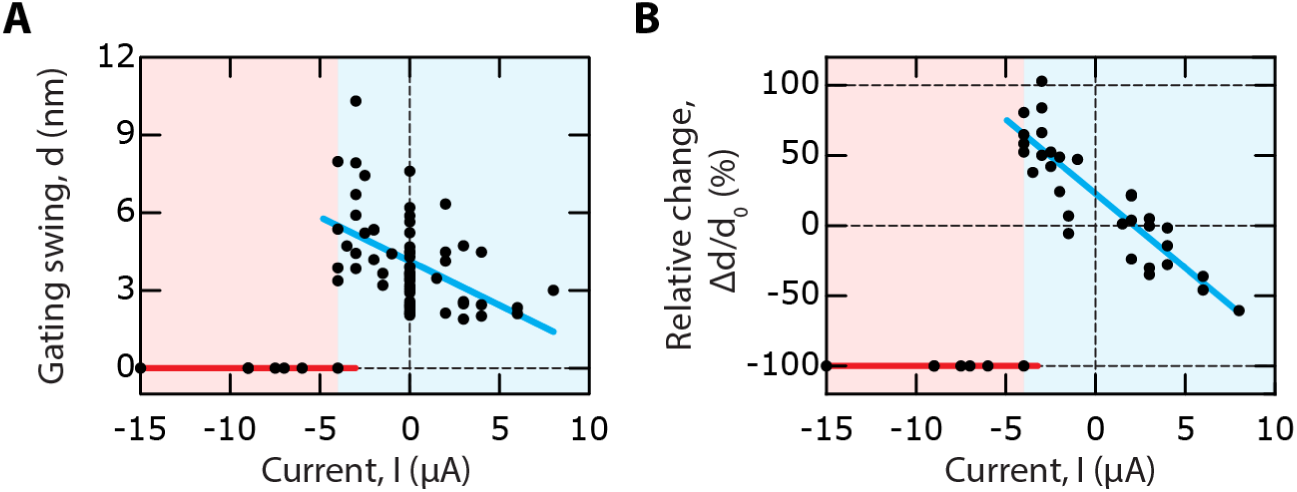
Electrical control of the gating swing. (**A**) For *I* ≥ −5 μA (cyan area), the gating swing, *d*, showed a negative correlation to the transepithelial current, *I* (Pearson correlation *r* = −0.61; p-value: 1 ⋅ 10^−7^; *n* = 63). Linear regression (cyan line): d [nm] = −0.3 · *I* [µ*A*] + 3.3 (*R*^2^ = 0.36). For I < −5 μA (red area), all the force displacement relations were linear: the gating swing had dropped to undetectable levels (red line). Under control conditions (*I* = 0), the gating swing had a mean value *d̅*_0_ = 3.7 ± 1.4 nm (mean ± SD, *n* = 29). (**B**) For I ≥ −5 μA (cyan area) the relative change, Δ*d*/*d*_0_, of the gating swing where Δ*d* = *d*(*I*) − *d*(*I* = 0) and *d*_0_ = *d*(*I* = 0) is plotted as a function of the transepithelial current, *I*. For *I* ≥ −5 μA, the relative change showed a strong negative correlation to the current (Pearson correlation *r* = −0.89; p-value: 3 ⋅ 10^−11^; *n* = 34). Linear regression (cyan line): Δ*d*/*d*_0_ [%] = −11 · *I* [µA] + 22 (*R*^2^ = 0.78). For I < −5 μA (red area), all the force displacement relations were linear: Δ*d*/*d*_0_ = −100% (red line). Same dataset as that used in Figure 3. After each measurement of a bundle’s force-displacement relation, the gating swing was estimated as *d* = *γF*_*G*_/(*K* − *K*_*SP*_), where *F*_*G*_ and *K* are respectively the gating force and linear stiffness resulting from a fit of the force-displacement relation with Equation 1 and *K*_*SP*_is the mean stiffness measured over an ensemble of hair bundles after disruption of their tip links. The projection factor *γ* = 0.14 was used to estimate the gating swing along the oblique axis of the tip link.

### 2.3 The transition between states of strong and weak gating force shows hysteresis

To further explore the transition between states of strong and weak gating force (Figure 2D and Figure 3C), we applied ramps of transepithelial current (**Figure 5A**) while the hair bundle was subjected to triangular mechanical stimulation (Figure 5B). This stimulation protocol allowed to monitor, continuously for the same hair bundle, how the mechanical properties varied as a function of the transepithelial current. The ramps were slow enough that the transepithelial current varied by only ∼0.2 µA over the time required to measure a force-displacement cycle (see Experimental Section). Because this variation was too small to elicit a significant variation of the gating force (Figure 3D), the mechanical measurements were quasi static. The mechanical stimulus elicited a force-displacement cycle with clockwise circulation, corresponding to energy dissipation.^[8]^ The cycle displayed a region of lower slope—gating compliance—on the positive and negative half cycles (Figure 5C), where the channels opened and closed, respectively. In addition, the height of the force-displacement cycle was larger within the region of channel gating. This property was previously shown to result from the contribution of the transduction channels’ gating force to hair-bundle friction, a phenomenon called gating friction, and the displacement at which the cycle had a maximal height (abscissa of disks in Figure 5C) was identified as the set-point deflection X_0_.^[8]^ In the following, the maximal half-height of the cycle, *ϕ*_*MAX*_, was used as a reporter for the magnitude of the gating force.

**Figure 5:**
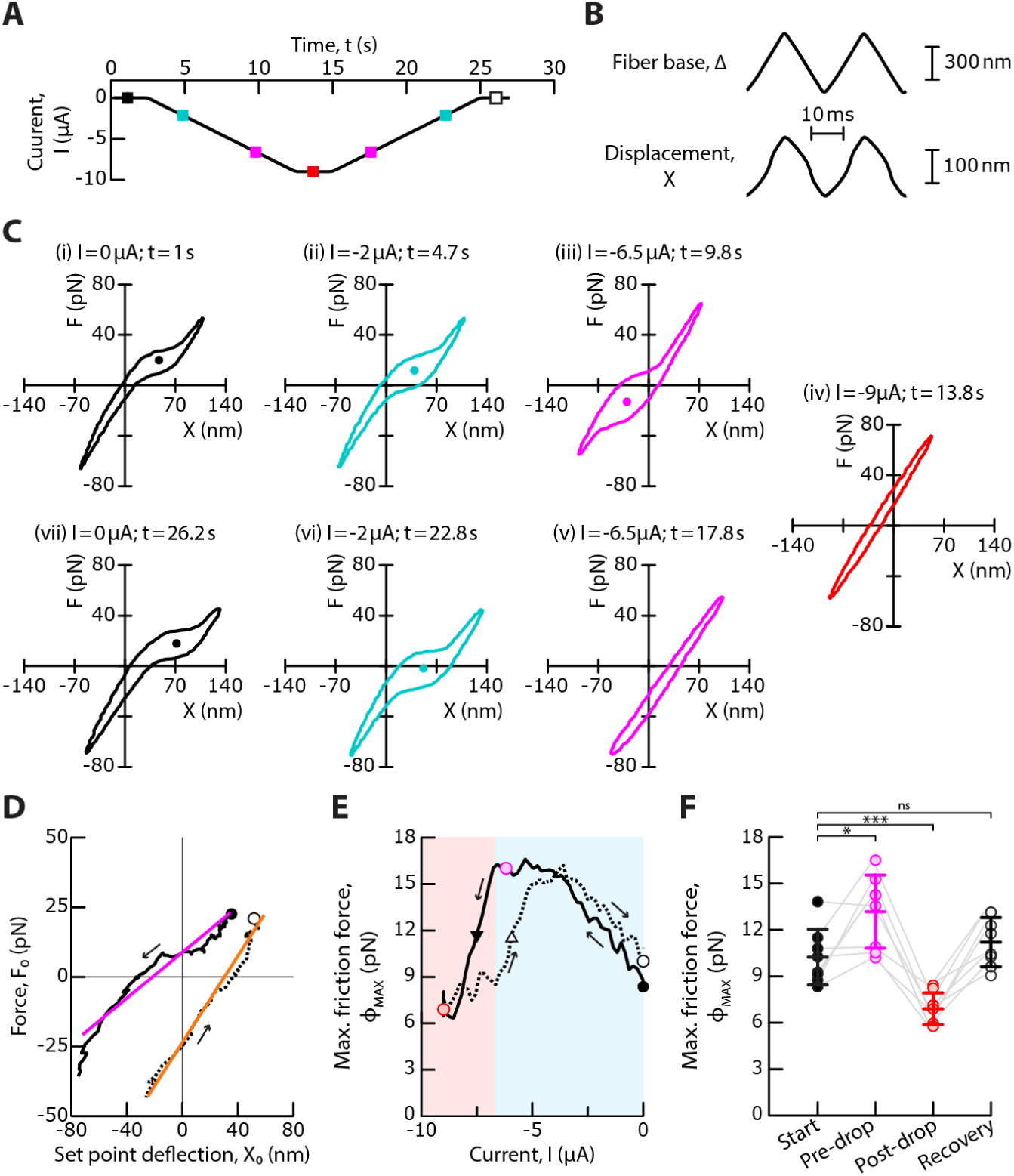
Transition between states of strong and weak gating force. **(A)** A descending ramp of a negative transepithelial current was followed by an ascending ramp of current at the same absolute rate of 0.9 μA ⋅ s^−1^. The colored square symbols indicate the time points of the analysis shown in (C). **(B)** The hair bundle was continuously moved back and forth (bottom) by imposing a periodic triangular movement to the stimulus fiber’s base, here with a peak-to-peak amplitude of 600 nm and at a frequency of 40 Hz (top). The movements were averaged over 10 stimulus cycles. **(C)** Force-displacement cycles for the values of the transepithelial current and at times indicated at the top of each sub-panel (see colored square symbols in (A)). **(D)** Trajectory of the point of inversion symmetry (X_0_, *F*_0_) in a force-displacement cycle, marked by disks in (C). Linear regressions to these trajectories: *F*_0_ [pN] = 0.41 · X_0_ [nm] + 9 (R^2^ = 0.92) (magenta) and *F*_0_ [pN] = 0.78 · X_0_ [nm] + 24 (R^2^ = 0.98) (orange), respectively. **(E)** Maximal friction force, *ϕ*_*MAX*_, as a function of the transepithelial current, *I*. Blue and red areas delineate regions of strong and weak gating force, respectively. Disks mark the value of *ϕ*_*MAX*_ at the start of the experiment (*I* = 0 µA, black), before (*I* = *I*_*DROP*_ + 1 μA with *I*_*DROP*_ = −6.9 µ*A*, magenta) and after (*I* = −9 μA, red) the drop of *ϕ*_*MAX*_, and at the end of the experiment (*I* = 0 μA, white). Black and white triangles indicate, respectively, the transition from a regime of strong to a regime of weak gating force and its reversal, at half the vertical distance between the magenta and red disks; their abscissas are shifted by 1.7 µA. **(F)** The distributions of *ϕ*_*MAX*_are shown for an ensemble of 8 hair bundles, at the start of each experiment (*I* = 0, black), before (*I* = *I*_*DROP*_ + 1 μA, magenta) and after (*I* = −9 μA, red) the drop of *ϕ*_*MAX*_, and at the end of the experiment (*I* = 0 μA, white). The hair bundle transitioned from a state of strong gating force to a state of weak gating force at more negative values of the current than a threshold *I* = *I*_*DROP*_ = −3.7 ± 1 μA (mean ± SD, *n* = 8). Mean values are shown as horizontal segments and error bars correspond to one standard deviation from the mean. Statistical significance was assessed using a paired t-test with p-values from left to right: 2 · 10^−2^, 8 · 10^−4^, 10^−1^. ns > 0.05, * p-value ≤ 0.05, *** p-value ≤ 0.001. In (D-E), black solid and dashed lines correspond to the descending and ascending ramps of transepithelial currents, respectively. Arrows indicate the direction of circulation along the curve from start (black disk) to finish (white disk).

As the transepithelial current decreased from zero to negative values, the region of gating compliance and gating friction in the force-displacement cycle shifted toward more negative displacements, corresponding to a decrease of the set-point deflection and thus opening of the transduction channels (Figure 5C, movie S1, Supplementary Information). Following the point of inversion symmetry in a force-displacement cycle resulted in a nearly linear trajectory of slope 0.4 ± 0.2 pN ⋅ nm^−1^ (*n* = 8; Figure 5D). In addition, the height of the force-displacement cycle, and thus the maximal friction force *ϕ*_*MAX*_ increased before saturating (Figure 5E). When the absolute magnitude of the negative current became large enough, corresponding to *I* ≤ *I*_*C*_ = −3.7 ± 1 μA and an endolymphatic potential *U* ≤ *U*_*C*_ ≃ −37 ± 10 mV (*n* = 8), the friction force *ϕ*_MAX_ dropped (Figure 5E, F and movie S1, Supplementary Information) until the nonlinearity vanished (Figure 5C, red). These observations on single hair cells (solid line in Figure 5E) mirrored the measured changes of the gating force upon application of a static transepithelial current in an ensemble of cells (Figure 3C). We note that gating friction generally depends on the activation time of the transduction channels, not just on the gating force.^[8]^ Detailed analysis of the force-displacement cycles suggests that the channel-activation time had increased by about 50%, on average, when the friction force *ϕ*_MAX_reached a saturating value upon application of a negative transepithelial current (Figure S3, Supplementary Information). The observed variation of hair-bundle friction (Figure 5E,F) thus resulted from modulation of both the channel-activation time and the gating force (Figure 3C). The polarity of the current ramp was then reversed. As the negative current increased, the set-point deflection increased and the point of inversion symmetry in the force-displacement cycle showed a nearly linear trajectory of slope 0.6 ± 0.2 pN ⋅ nm^−1^ (*n* = 8; Figure 5C and D). In addition, the signatures of gating compliance and gating friction, associated to nonlinear force-displacement relations, were retrieved. However, as shown respectively in subpanels (v) and (iii) of Figure 5C, the gating force could remain weak on the ascending ramp at current values where the gating force was strong on the descending ramp. Accordingly, the relation between the friction force *ϕ*_*MAX*_ and the transepithelial current *I* displayed hysteresis (Figure 5E). Over an ensemble of *n* = 8 cells, the transition from a regime of strong to a regime of weak gating force and its reversal were shifted by 2.3 ± 0.8 μA, corresponding to a difference of about 20 mV difference in endolymphatic potential.

### 2.4 The transition kinetics between the two states occurs on a sub-second timescale

To characterize the kinetics of the transition between states of strong and weak gating force, we applied large negative steps of transepithelial current, here *I* = −8 μA (**Figure 6**), while continuously measuring a hair bundle’s force-displacement cycle in response to triangular mechanical stimulation. As expected from the effects of current ramps (Figure 5), the force-displacement cycle shifted in the negative direction along the displacement axis and the force-displacement cycle lost the signatures of gating compliance and gating friction (Figure 6A, movie S2, Supplementary Information). The time courses of the average hair-bundle position on the cycle, denoted 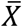 in the following, and of the maximal friction force *ϕ*_*MAX*_ were quantified by measuring the 10 − 90% decay and rise times following the onset and offset of the current step, respectively. Over an ensemble of *n* = 4 hair cells, the characteristic time, 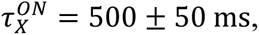 of negative hair-bundle motion evoked by the current step was fivefold longer than that, 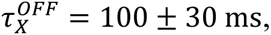 necessary to return to the initial position after the current was turned off (Figure 6B). The friction force *ϕ*_*MAX*_ showed an opposite behavior: its decay time, 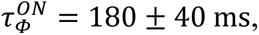 after the onset of the current step was twofold shorter than its rise time, 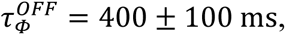 during recovery (Figure 6B). Note that the time required to measure a force-displacement cycle was here of only 20 ms. This duration was about ten times shorter than 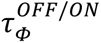 and thus short enough to ensure a quasistatic assessment of hair-bundle mechanics (Experimental Section).

**Figure 6:**
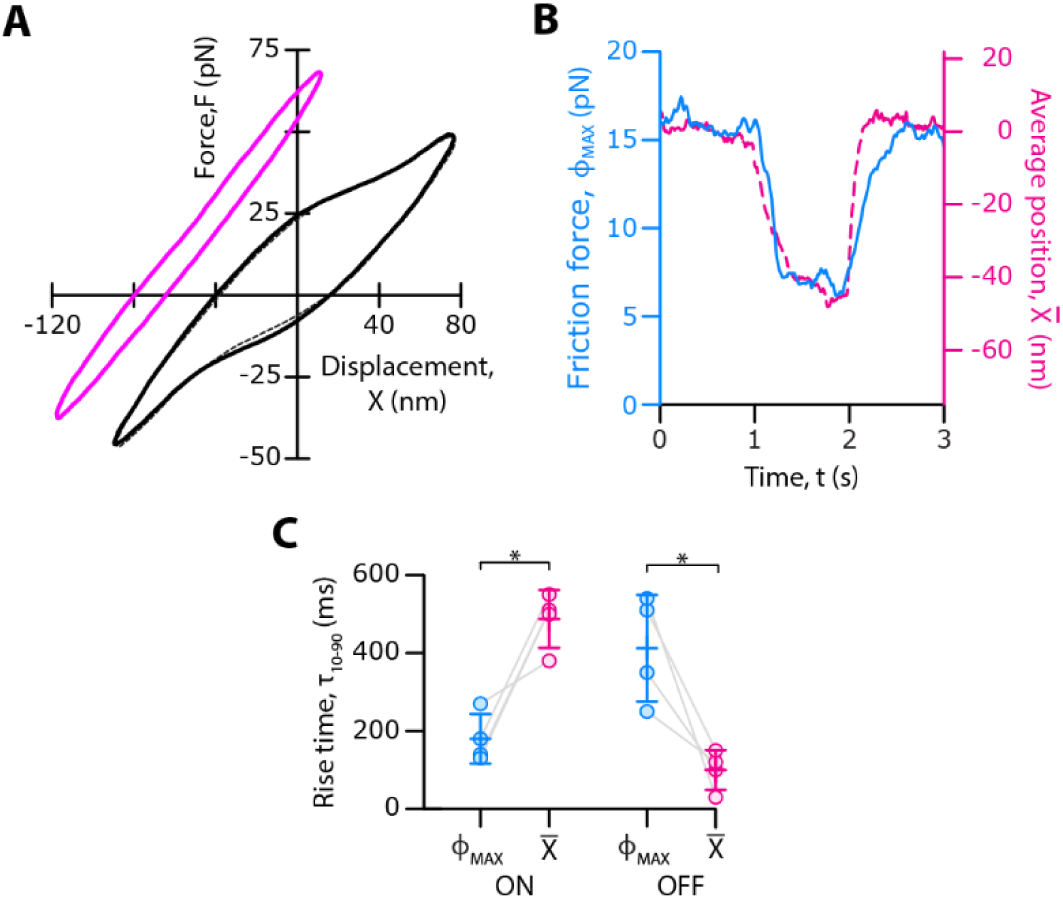
Transition kinetics between states of strong and weak gating force. **(A)** Force-displacement cycle here averaged over 50 cycles of stimulation, under control conditions (*I* = 0; black solid line), 400 ms after the onset of a step of transepithelial current (*I* = −8 µA; magenta) and 600 ms after termination of the current step (*I* = 0; black dashed line). **(B)** Time course of the maximal friction force, *ϕ*_*MAX*_, (blue line) and of the average displacement, 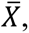 (dashed magenta line) in a force-displacement cycle. The current step started at *t* = 1 s and ended at *t* = 2 s. **(C)** For all the 4 different hair bundles that followed the same protocol, time, *τ*_10−90_, required for the maximal friction force, *ϕ*_*MAX*_(blue disks), and the average displacement, 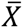 (magenta disks), to transition from 10% to 90% of the total change after the onset (ON) and the termination (OFF) of a step of transepithelial current (*I* = −8 µA). Mean values are shown as horizontal segments and error bars correspond to one standard deviation from the mean. Statistical significance was assessed using a paired t-test with p-values from left to right: 2 · 10^−2^, 3 · 10^−2^; * p-value ≤ 0.05. The base of the stimulus fiber followed a periodic triangular waveform of motion, with a peak-to-peak amplitude of 600 nm and at a frequency of 100 Hz. For the data shown in (B) and (C), the force-displacement cycles were averaged over only two cycles of stimulation.

There was thus a separation of timescales between the changes of 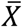 and *ϕ*_*MAX*_. At the onset of the current step, gating compliance vanished before the negative shift had completed: 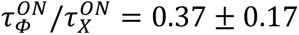 (Figure 6B,C). In contrast, after the external transepithelial current was turned off, the mechanical nonlinearity recovered more slowly than the average position of the hair bundle: 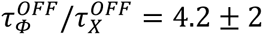 (Figure 6B,C).

### 2.5 Reducing the endolymphatic Ca^2+^ concentration replicates the effects of a negative endolymphatic potential

A negative endolymphatic potential may have mediated its effects on the gating force by reducing the magnitude of the Ca^2+^ influx through the transduction channels. To test this hypothesis, we used iontophoresis of a Ca^2+^ chelator to reduce the endolymphatic Ca^2+^ concentration near a hair bundle and, in turn, the Ca^2+^ influx through the channels under resting conditions. In response to descending and ascending ramps of iontophoretic current (**Figure 7A**), or equivalently of Ca^2+^ concentration, we again observed an increase in hair-bundle friction followed by a reversible but hysteretic transition between states of strong and weak gating force (Figure 7B,C, movie S3, Supplementary Information). Repeating the experiments over an ensemble of *n* = 8 cells confirmed the robustness of these effects (Figure 7D) and how they mirrored those observed upon modulation of the endolymphatic potential (Figure 5). In addition, the force-displacement relation of a hair bundle could show strong gating compliance under control conditions but reversibly become linear upon step application of a Ca^2+^ chelator, with no noticeable change in the linear stiffness of the hair bundle (Figure 7E). The same behavior was observed in *n* = 7 hair cells. Thus, enough calcium in endolymph was required for the hair bundle to evince strong gating forces.

**Figure 7:**
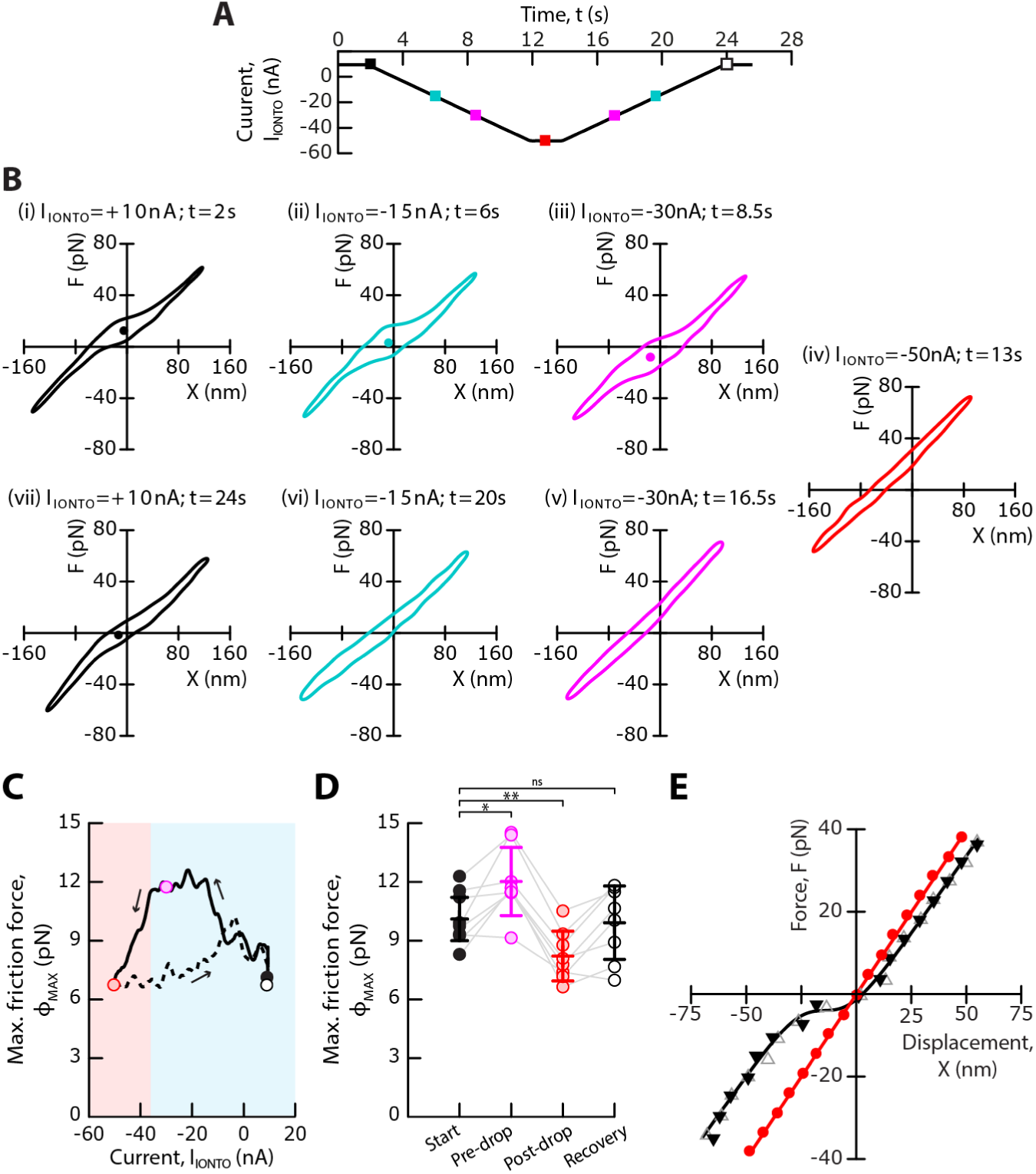
Replicating the effects of negative endolymphatic potentials with iontophoresis of a Ca^2+^ chelator. **(A)** We used iontophoresis of a calcium chelator to decrease the extracellular Ca^2+^ concentration in the vicinity of a hair bundle. A descending ramp of a negative iontophoretic current, *I*_IONTO_, which was carried in part by a Ca^2+^chelator, was followed by an ascending ramp of current at the same absolute rate of 6.6 nA ⋅ s^−1^. The colored square symbols indicate the time points of the analysis shown in (B). **(B)** Each sub-panel shows the bundle’s force-displacement cycle in response to a triangular mechanical stimulus for the value of the iontophoretic current, *I*_*IONTO*_, and at the time, *t*, indicated at the top. **(C)**. Maximal friction force, *ϕ*_*MAX*_, as a function of the iontophoretic current, *I*_*IONTO*_. Blue and red areas delineate regions of strong and weak gating force, respectively. Disks mark the value of *ϕ*_*MAX*_ at the start of the experiment (*I*_*IONTO*_ = +10 nA, black), before (*I*_*IONTO*_ = *I*_*DROP*_ + 5 nA with *I*_*DROP*_ = −35 nA, magenta) and after (*I*_*IONTO*_ = −50 nA, red) the drop of *ϕ*_*MAX*_, and at the end of the experiment (*I*_*IONTO*_ = +10 nA, white). Arrows indicate the direction of circulation along the curve. Solid and dashed lines correspond to the descending and ascending ramps of iontophoretic currents, respectively. **(D)** The distributions of *ϕ*_*MAX*_are shown for an ensemble of 8 hair bundles, at the start of the experiment (*I*_*IONTO*_ = +10 nA, black), before (*I*_*IONTO*_ = *I*_*DROP*_ + 5 nA, magenta) and after (*I*_*IONTO*_ = −50 nA, red) the drop of *ϕ*_*MAX*_, and at the end of the experiment (*I*_*IONTO*_ = +10 nA, white). The hair bundle transitioned from a state of strong gating force to a state of weak gating force at more negative values of the current than a threshold *I*_*IONTO*_ = *I*_*DROP*_ = −40 ± 10 nA (mean ± SD, *n* = 8). Mean values are shown as horizontal segments and error bars correspond to one standard deviation from the mean. Statistical significance was assessed using a paired t-test with p-values from left to right: 1.5 · 10^−2^, 5 · 10^−3^, 6 · 10^−1^; ns > 0.05; * p-value ≤ 0.05, ** p-value ≤ 0.01. (**E**) The force-displacement relation of a different hair bundle displays a region of gating compliance under control conditions (black triangles) but was linear in the presence of a steady iontophoretic current of −60 nA (red circle). This effect was reversible (grey triangles). Each force-displacement relation was fitted by Equation 1 (solid lines; parameter values in Table S1). In (B-D), a periodic triangular waveform of motion was applied to the base of the stimulus fiber, with a peak-to-peak amplitude of 600 nm and at a frequency of 40 Hz. In (E), force-displacement relations were instead measured in response to step displacements at the base of the stimulus fiber (see Experimental Section).

## 3. Conclusion

In this work, we showed that varying the endolymphatic potential reversibly modulates the gating force, thus serving as a control parameter for the mechanosensitivity of the hair cell to minute deflections of its hair bundle. This bioelectric effect is large. The gating force increased continuously by up to twofold upon application of negative potentials of moderate magnitude (0 ≥ *U* ≥ −50 mV) and conversely, could be reduced by up to 80% of the control value at positive potentials (Figure 3D). A major finding of our study is that the gating force varied because the gating swing varied, with only minor effects on the gating-spring stiffness (Figures 3D and 4).

In addition, the hair cell switched from a state of strong gating force to a state of weak gating force when the endolymphatic potential became more negative than a critical value *U* = *U*_*C*_ ≃ −50 mV (Figures 2D, 3C, 5C and 4E). This observation revealed an unexpected transition between two functional states of the hair bundle—one with low mechanosensitivity and one with high mechanosensitivity: a quiescent state of undetectable gating force in hair-bundle mechanics, which is suggestive of gating swings of about one nanometer or less, and an oscillatory state of strong gating force with gating swings of several nanometers, up to 10 nm. At such negative potentials (*U* ≤ *U*_*C*_), the electromotive force driving the influx of cations through the transduction channels is expected to be greatly reduced. Because reducing the Ca^2+^ concentration near a hair bundle evoked a similar transition (Figure 7), the strong gating forces and large gating swings associated to high mechanosensitivity likely required a resting Ca^2+^ influx through the transduction channels that was large enough.

Altogether, the variation of the transduction channels’ gating force and the gating swing as a function of the endolymphatic potential was nonmonotonic, indicating that mechanosensitivity is maximal within a limited range of potential and, in turn, of Ca^2+^ influx under resting conditions (**Figure 8**).

**Figure 8:**
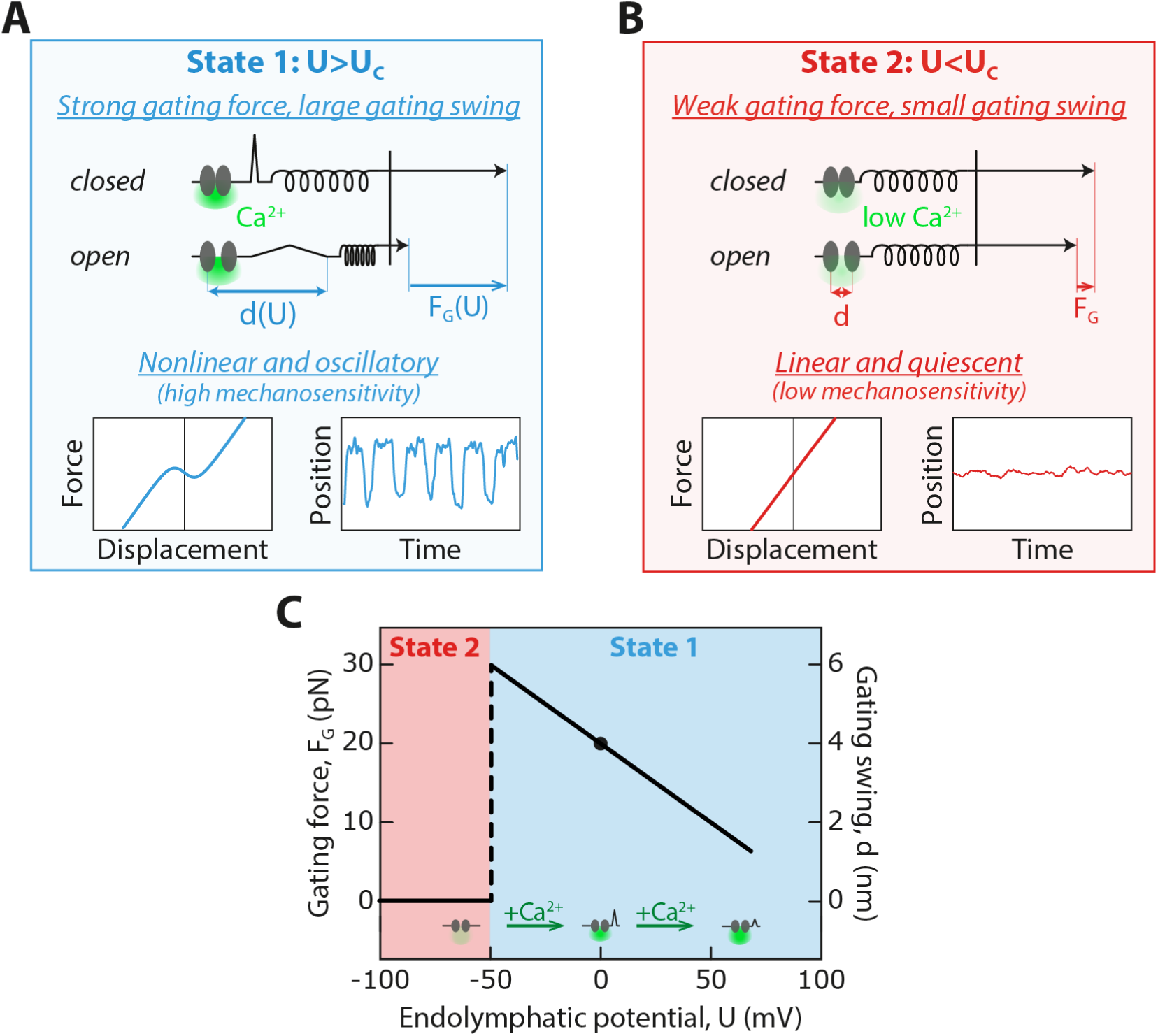
Conceptual model of electrical gating-force and gating-swing control via intracellular calcium. (**A**) By controlling the electromotive force that drives Ca^2+^ entry, the endolymphatic potential, *U*, controls the time-averaged intracellular Ca^2+^ concentration near the channels’ pore (green halo) as the channels fluctuate between their open and closed states under resting conditions. For endolymphatic potentials, *U* ≥ *U*_*C*_, larger than a critical value, *U*_*C*_ ≃ −50 *mV*, the intracellular environment is relatively rich in Ca^2+^, which allows the transduction channels to operate in a state of strong gating force, *F*_*G*_(*U*), and large gating swing, *d*(*U*). A movement within the transduction-channel protein complex must magnify the small, nanometer-sized, conformational change associated to opening and closing of the channels’ pore. In this state—State 1, the hair bundle shows a nonlinear region of strong gating compliance in its force-displacement relation and, in turn, spontaneous oscillations. These active movements have been previously shown to foster frequency-selective amplification of mechanosensitivity to weak periodic stimuli. (**B**) For endolymphatic potentials, *U* ≤ *U*_*C*_, the intracellular ionic environment of the channel is poor in Ca^2+^. Consequently, the transduction channels operate in a state of weak gating force and small gating swing—State 2. In this state, the hair bundle is quiescent and endowed with a linear force-displacement relation: there is no detectable signature of channel gating in hair-bundle mechanics, which suggests that the gating swing is less than about one nanometer. Mechanosensitivity in State 2 is much smaller than that in State 1. (**C**) In State 1 (cyan area), the endolymphatic potential, *U*, continuously and reversibly modulates the magnitude, *F*_*G*_, of the gating force exerted by the transduction channels on the whole hair bundle (left axis) by changing the channels’ gating swing, *d* (right axis). There is a transition between State 1 and State 2 (red area) at a value of the endolymphatic potential *U* = *U*_*C*_ ≃ −50 *mV*. Modulation of the intracellular Ca^2+^ concentration at rest by the endolymphatic potential and its effect on the gating swing is shown schematically at the bottom. Our work suggests that enough intracellular Ca^2+^ is required to prompt large gating forces (*U* ≥ *U*_*C*_; State 1) but that further Ca^2+^ increase results in a continuous decrease of the gating force and gating swing. Under control conditions (*U* = 0, black disks), the transduction channels operate in a state of strong gating force.

### 3.1 Physiological role of the endolymphatic potential

Like other epithelia, the sensory epithelia that house the hair cells in the vertebrate inner ear are polar tissues that experience a transepithelial potential under natural conditions.^[34]^ This potential generally lies within a range 1 − 10 mV, except in the mammalian cochlea where the endolymphatic potential can reach +100 mV. Remarkably enough, this high value of the endolymphatic potential is associated with a Ca^2+^ concentration in endolymph, about 20 µM, which is about one of order of magnitude lower than in hearing organs from non-mammalian species or than in vestibular organs. Our results on frog saccular hair cells (Figures 3 and 7) raise the possibility that the combination of a large endolymphatic potential and a low Ca^2+^ concentration in the cochlea might maintain a Ca^2+^ influx within the optimal range to allow for large gating forces while boosting the magnitude of transduction currents. In support of this inference, a mechanical nonlinearity attributed to hair-bundle mechanics has been measured in an ex-vivo preparation of the Gerbil cochlea, but only under conditions that replicate a normal endolymphatic potential and Ca^2+^ concentration.^[13,35]^ This nonlinearity disappears when no endolymphatic potential is applied, which makes sense within the framework of our results: the potential is then effectively very negative with respect to natural conditions—by 100 mV, perhaps enforcing a similar state of weak gating force as we report here in frog upon application of a potential that is negative enough (Figure 2D, 3C and 5). Note that immersing the sensory epithelium in artificial perilymph, as is routinely performed in the field, is also expected to lead to weak gating forces but here as a result of a too high Ca^2+^ influx. In the cochlea, the Ca^2+^ concentration in perilymph is indeed nearly two orders of magnitude higher than in endolymph. Our results on frog saccular hair cells thus help putting ex-vivo measurements on mammalian hair cells into perspective. In addition, our observations also imply that a spatial gradient in the endolymphatic potential, as observed in vivo in the mammalian cochlea,^[36]^ might provide a mechanism to tune the gating force and thus mechanosensitivity across hair cells.

In our experiments on hair cells from the frog saccule, control conditions approach natural conditions: the top and bottom compartments have ionic compositions that mimic those of endolymph and perilymph, respectively, and there is no transepithelial potential. Also, our approach is based on force measurements, which in contrast to standard electrophysiological approaches does not disrupt the integrity of the hair cell or their internal ionic concentrations. Under these nearly natural conditions, the gating force is strong, as a matter of fact strong enough to evoke negative stiffness, promote spontaneous oscillations of the hair bundle and mechanical amplification of weak periodic stimuli.^[9,23,24]^ We applied endolymphatic potentials within a range of ±100 mV. While this range departs from natural conditions, pushing the hair cells into uncharted territory disclosed gating-force control by the endolymphatic potential and also indicated that the hair cell operates near an optimum of mechanosensitivity under natural conditions.

It is worth noting that the difference in electrical potential between endolymph and the hair-cell interior likely constitutes the relevant control parameter, not the endolymph potential per se. From this perspective, modifying the endolymphatic potential may serve as a proxy for evaluating the effects of varying the hair cell’s transmembrane potential—at least insofar as mechanosensitivity of the transduction channels is concerned. The efferent nervous system is known to provide descending feedback from the brain to tune sensitivity of the auditory system.^[37]^ Activation of the efferent system leads to hyperpolarization of the hair cell.^[38–40]^ Assuming that hyperpolarization of the hair cell has similar effects on transduction channels’ operation as raising the endolymph potential to more positive values, our work opens the possibility that efferent activation might help desensitize hair cells by reducing the transduction channels’ gating force and gating swing (Figure 8). More generally, any perturbation evoking a sustained change of the endolymphatic potential, of the intracellular potential, or of the intracellular calcium concentration near the channels’ pore would be expected to affect the gating force and, in turn, hair-cell mechanosensitivity.

### 3.2 Effects of Ca^2+^ on hair-bundle mechanics

Varying the endolymphatic potential afforded a means to reversibly and robustly control the electric field in the membrane near the transduction channels and in turn the electromotive force that controls the magnitude of transduction currents, including the Ca^2+^ influx into the hair cell. Reducing the Ca^2+^ concentration near a hair bundle resulted in similar effects on gating compliance and gating friction than applying negative endolymphatic potentials (Figure 7). This finding indicates that negative endolymphatic potentials modulated the gating force, at least in part, by reducing the Ca^2+^ influx through the transduction channels, whereas positive potentials do the opposite. We cannot exclude, however, that the endolymphatic potential also directly affected channel gating by mediating an electric force on the transduction-channel protein complex. Depolarization of the hair cell has been previously shown to evoke fast movements of the hair bundle, even with channels blocked and thus with no change in the Ca^2+^ influx though the channels, which may be indicative of a rearrangement of the channel protein complex.^[41,42]^

Calcium has previously been shown to affect the channel activation time, the operating point of the transducer and the kinetics of the active adaptation process that drives active hair-bundle motility.^[4,24,26,27,30,31,43,44]^ Moreover, there is some evidence that the extracellular calcium concentration can modulate the stiffness of the hair bundle.^[27,33,45]^ In our experiments, negative endolymphatic potentials produced effects on spontaneous hair-bundle oscillations (Figure S2, Supplementary Information) that were qualitatively similar as those reported earlier in response to depolarization of the hair cell ^[41,46]^ or to a decrease of the Ca^2+^ concentration in endolymph.^[26]^ Conversely, positive endolymphatic potentials produced similar effects as hyperpolarization of the hair cell or increasing the endolymphatic Ca^2+^ concentration.

Our findings demonstrate that calcium does more than modulate the operating point of the transducer and the kinetics of channel gating and adaptation: it also affects the magnitude of the gating force. Previous models had postulated stiffer gating springs at lower Ca^2+^,^[26,47]^ in apparent contradiction with recent in-vitro measurements of PCDH15 elasticity.^[48]^ Our observations (Figures 3, 4 and 7) suggest instead that the effect of Ca^2+^ on gating-spring stiffness is small compared to that on the gating swing, at least under our experimental conditions.

### 3.3 Implications for the mechanism of channel gating

As recognized earlier,^[9]^ we observed that the gating swing (Figure 4) can be nearly one order of magnitude larger than the expected diameter of the transduction channels’ pore, which has been estimated at about 1.2 nm.^[2]^ Hair cells thus appear to have implemented a mechanism to magnify the conformational change associated to opening and closing of the transduction channel’s pore, a feature also relevant to other mechanosensitive channels.^[49]^ In hair cells, the ion channels that mediate mechanoelectrical transduction are part of a molecular complex that includes TMC1—the pore-forming subunit,^[50]^ TMIE, CIB2, LHFPL5 and the lower end of the tip link, PCDH15.^[51,52]^ In addition, the complex may comprise two copies of each protein, forming a dimeric assembly with an axis of twofold symmetry ^[53,54]^ and yielding a total of at least 10 proteins.

There is currently no evidence for a large conformational change in a single component of the transduction complex (e.g. in TMC1, see ^[55]^). However, interactions between the multiple components of the transduction apparatus might lead to a collective rearrangement upon opening of the channel’s pore, effectively producing a large gating swing. In this view, the gating swing is not the result of a single conformational change in a single protein—say pore opening in TMC1, but instead of an avalanche of conformational changes in multiple proteins operating in series. In addition, the lipid membrane that houses the transduction-channel protein complex may play a role in the implementation of this effective lever arm on pore opening. Channel gating has been proposed to elicit a membrane-mediated interaction between two channel complexes, each attached to one strand of the PCDH15 dimer, leading to a large movement within the membrane that magnifies the apparent gating swing.^[56]^ Alternatively, as proposed for eukaryotic Piezo channels based on structural evidence,^[49]^ the transduction-channel protein complex may not sit in a flat lipid membrane but instead curve the membrane. A change in membrane deformation upon opening of the channel’s pore might then magnify the gating swing.

Whatever the molecular substrate for the gating swing, our work indicates that the transepithelial potential and the extracellular calcium concentration can continuously modulate the magnitude of the movement associated to channel gating when the hair bundle operates in a state of strong gating force and large gating swing (State 1 in Figure 8). In addition, under conditions for which the calcium influx through the channels becomes too low (State 2 in Figure 8), we found that the gating force drops to undetectable levels. The transduction channels have been shown to retain their ability to open and close even when hair cells are depolarized to extreme values, up to +100 mV, ^[57–59]^ which, like the very negative endolymphatic potentials applied here, strongly opposes Ca^2+^ entry into the hair cells. In our experiments, the observed transition to a state of weak gating force at endolymphatic potentials *U* ≤ *U*_*C*_ is therefore likely attributable to a drop in the magnitude of the gating swing rather than to a loss of the channels’ ability to gate. Our observations suggest that the channel-protein complex must be primed by enough intracellular calcium to elicit a large gating swing.

What could be the mechanisms by which Ca^2+^ modulates the gating swing? Recent evidence opens two attractive possibilities. First, calcium was shown to evoke the dissociation of a cytosolic region of TMC1 from CIB2—the calcium- and integrin-binding protein.^[60,61]^ By promoting a tight interaction between CIB2 and TMC1, low calcium might prevent large gating swings upon opening of the channel’s pore and, conversely, that enough intracellular Ca^2+^ is necessary to promote a state of high mechanosensitivity of the transduction channel. Hysteresis upon a continuous back-and-forth variation of the endolymphatic potential (Figure 5) or of the extracellular concentration (Figure 7), and the relatively short timescale of the transition between states of strong and weak gating force (Figure 6) provide further support for the bistability of the channel-protein complex. Calcium modulation of the interaction between CIB2 and TMC1 might contribute to the switch-like behavior of the gating swing (Figure. 8).

Second, calcium may also modulate the effective size of the gating swing by affecting the mechanical properties of the lipid membrane that houses the transduction channels.^[56,62]^ Diffusivity of the lipids in the membrane of stereocilia varies with calcium and is modulated by the presence of functional transduction channels.^[62,63]^ These observations are consistent with a dual role of the transduction channels as both ion channels and membrane scramblases,^[64]^ which mediate passive bidirectional movement of lipids between the two leaflets of a biological membrane. Modulation of membrane composition is expected to affect channel gating.^[65]^ In addition, should the membrane play a role in force transmission from the tip link to the channel—a departure from the more widely accepted tether model of channel activation, one would expect high-pass filtering of external forces and, reciprocally, of the mechanical correlate of channel gating. This regulatory mechanism can only lead to underestimates but cannot, by itself, explain the large gating swings reported here. Altogether, calcium-mediated modulation of membrane properties may contribute to the modulability of the gating force and gating swing. To cope with the physical requirements of hearing, hair cells have evolved the ability to respond to sound stimuli that vary over timescales as short as a few tens of microseconds, detect deflections of their hair bundle as small as one nanometer or less, and mediate sharp frequency selectivity over three orders of magnitude of sound frequencies.^[66]^ These unique capabilities rely on a transduction-channel protein complex with a highly intricate molecular composition and architecture.^[52,54]^ Inasmuch that the molecular underpinnings of mechanoelectrical transduction by hair cells remains similar across species and organs, our force measurements on hair bundles from frog saccular hair cells demonstrate that the effective conformational change associated to channel gating—the gating swing—can be modulated over a broad range by varying physiological control parameters. The modulability of the gating swing provides a mechanism to tune passive and active mechanosensitivity of the hair cell while maintaining the molecular composition of the transduction channels. Elucidating the molecular rearrangement associated to channel gating and how it can be modulated remain major challenges for future studies.

## 4. Experimental Section

### Experimental preparation

All experimental procedures were approved by the Ethics committee on animal experimentation of the Institut Curie; they complied with the European and French National Regulation for the Protection of Vertebrate Animals used for Experimental and other Scientific Purposes (Directive 2010/63; French Decree 2013–118). Details of the experimental preparation have been published elsewhere.^[26,27]^ Briefly, an excised preparation of the frog’s saccule (Rivan 92 ^[67]^) was glued with tissue-compatible surgical glue (Histoacryl, B. Braun) over a ∼1 mm hole in a plastic disk (Figure 1A). The preparation was then secured in a two-compartment chamber, which exposed the apical and basal aspects of the sensory tissue to fluids with different ionic compositions to mimic physiological conditions. The basal bodies of the hair cells were bathed in artificial perilymph, a standard saline solution containing (in mM): 110 NaCl, 2 KCl, 1.5 CaCl_2_, 3 D-glucose and 5 Hepes. Hair bundles instead projected into artificial endolymph with (in mM): 2 NaCl, 118 KCl, 0.25 CaCl_2_, 3 D-glucose, and 5 Hepes. Each solution had a pH of ∼ 7.3 and an osmotic strength of ∼ 225 mmol ⋅ kg^−1^. To disconnect the hair bundles from the overlying otolithic membrane, the apical surface of the preparation was exposed for 20 min to endolymph supplemented with 67 μg ⋅ mL^−1^of the protease subtilisin (type XXIV, Sigma). The otolithic membrane was then peeled off to obtain access to individual hair bundles. Experiments were performed at a temperature of 20°C.

### Microscopic apparatus and mechanical stimulation

The preparation was viewed through a × 60 water-immersion objective of an upright microscope (BX51WI, Olympus). The mechanical properties of single hair bundles were characterized using published methods.^[8,27]^ In short, the tip of a stimulus fiber was affixed to the kinociliary bulb of an individual hair bundle and imaged at a magnification of × 944 onto a displacement monitor comprising a dual photodiode. Stimulus fibers were fabricated from borosilicate capillaries (TW120-3, World Precision Instrument) and coated with a thin layer of gold-palladium to enhance contrast (Hummer 6.1, Anatech ltd.). The fiber was secured by its base to a stack-type piezoelectric actuator (PA-8/14, Piezosystem Jena) driven by a custom-made power supply (Elbatech). Its stiffness, *k*_*F*_, was estimated from the analysis of thermal fluctuations at the fiber’s tip.

In a first stimulation protocol, a series of step displacements, Δ, was applied to the fiber’s base. The standard protocol consisted in the application of 7 to 9 successive pairs of step displacements in opposite directions with a magnitude growing with a constant increment, within the range 20 − 40 nm. The force, *F* = *k*_*F*_ (Δ − X), and the bundle’s displacement, X, were estimated 5 − 9 ms after the stimulus onset. This time was long enough for the open probability of the transduction channels to have reached a new steady state and for the frictional contribution to the applied force to have vanished, but short enough to limit the effect of adaptation. The resulting force-displacement relation, *F*(X), was fitted by the equation ^[5,24]^:

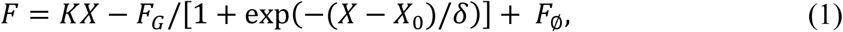

from which we estimated the bundle’s linear stiffness, *K*, the gating force, *F*_*G*_, for the whole hair bundle, and the set-point deflection, X_0_. The parameter *F*_∅_ ensured that *F* = 0 at X = 0. Because we did not use displacement-clamp feedback to stabilize positions of negative stiffness,^[9]^ the characteristic displacement, *δ*, could not be exploited. In this protocol, both photodiode and command signals were acquired and generated at a sampling rate of 2.5 kHz and the half-power frequency of the lowpass filter was fixed at 1 kHz. A force-displacement relationship was obtained by averaging 7 to 15 repetitions of the same stimulation protocol, which collectively took 10 to 20 seconds.

In a second stimulation protocol, we applied a symmetric triangular waveform of motion to the fiber’s base, at a frequency of either 40 Hz or 100 Hz and with a peak-to-peak amplitude within the range 600 − 1000 nm. The force applied to the hair bundle was estimated using a detailed description of the fiber’s vibrational modes.^[8]^ This protocol resulted in force-displacement cycles with counterclockwise circulation. The half-height at any position, X, in the cycle provides an estimate of the average friction force *ϕ*(X) = [*F*^+^(X) − *F*^−^(X)]/2 impeding the hair bundle motion at this position, where *F*^±^(X) represents the force in the positive and negative half cycle of stimulation, respectively. The friction force displayed a maximum, *ϕ*_MAX_ = *ϕ*(X_0_), at a displacement that was identified earlier as the set-point deflection X_0_in Equation 1.^[8]^ The force-displacement cycle also displayed a point of inversion symmetry of coordinates (X_0_, *F*_0_ = [*F*^+^(X_0_) + *F*^−^(X_0_)]/2). The shift, ΔX, between the positions where the hair-bundle speed reached a maximal value, |*d*X/*dt*| = *V*_*MAX*_, in the positive and negative half cycle of stimulation provided an estimate of the activation time of the transduction channels, *τ* = ΔX/(2*V*_*MAX*_).^[8]^ In this protocol, the force-displacement cycle was averaged over ten periods of the triangular stimulus at 40 Hz or, when a fast assessment of hair-bundle mechanics became necessary (as for the data shown in Figure 6), over only two periods of a 100-Hz stimulus. Triangular stimulation thus afforded a means to probe hair-bundle mechanics in a period of time of 20—250 ms, which was 40—1,000 times shorter than the time required to measure a force-displacement relation with the step-stimulation protocol described above. Both photodiode and command signals were acquired and generated at sampling rate of 25 kHz. The half-power frequency of the filter was fixed at 12.5 kHz for the photodiode signals and 0.5 kHz for the command signals.

### Control of the transepithelial current and potential

Using a current source (Linear Stimulator Isolator A395, World Precision Instruments) connected to a pair of silver chloride electrodes (Figure 1A), we applied steps of transepithelial current with a magnitude that was varied within a range of ±10 µA. In some preparations, we measured the transepithelial potential, *U*(*t*), evoked by a step of transepithelial current, *I*_*STEP*_ = 1 μA (Figure 1B), using an additional pair of electrodes connected to the headstage of a differential amplifier (voltage gain × 50; Model 3000, Systems Inc). The time course of the transepithelial potential was well described by an exponential relation *U*(*t*) = *U*_*MAX*_ exp(−*t*/*τ*), with a characteristic timescale τ = 230 ± 50 μs (mean ± SD, *n* =13). Over this ensemble of 13 saccules, we found a transepithelial resistance *R* = *U*_*MAX*_/*I*_*STEP*_ = 10 ± 2 kΩ and capacitance *C* = 23 ± 4 nF, which accords with previous estimates.^[22]^ Thus, over the range of transepithelial currents that we explored, the potential into the artificial endolymph that bathed the hair bundles—the endolymphatic potential— varied within a range of ±100 mV with respect to the potential in the perilymphatic compartment. In preparations for which we did not measure the transepithelial resistance, the transepithelial potential was estimated by multiplying the applied transepithelial current by the ensemble-averaged resistance given above, thus with a precision of 20%. We note that the tight junctions between hair cells and supporting cells, despite their names, constitute the least resistive element in the equivalent electrical circuit associated to the frog saccule and thus dictate the resistance of the epithelium.^[22]^ The effect of a transepithelial current on the Ca^2+^ and K^+^ influx in to the hair cells is thus thought to be indirect: the transepithelial current produces a transepithelial potential, which in turn affects the magnitude of the electromotive force that drives the ions through the transduction channels.

In other experiments, we applied successive ramps of descending and ascending transepithelial currents at an absolute rate within the range of 0.5 − 0.9 µA · s^−1^. These rates were low enough to ensure that the variation of the transepithelial current remained small during the time required for probing hair-bundle mechanics, allowing these mechanical measurements to remain quasistatic. The minimal current was maintained for 1 − 2.5 s before the ascending ramp started.

### Iontophoresis of a calcium chelator

We used iontophoresis of a calcium chelator to decrease the extracellular calcium concentration in the vicinity of individual hair bundles.^[15,27]^ Coarse microelectrodes were fabricated from borosilicate capillaries (TW120F-3, World Precision Instrument) with a pipette puller (P97, Sutter Instrument); their resistance ranged from 0.8 to 1.5 MΩ when filled with 3-M KCl and immersed in the same solution. The electrode’s tip was positioned at ∼3 μm from the hair bundle.

In a first series of experiments, the electrodes were filled with 400-µM pentasodium triphosphate. Triphosphate allowed to decrease the calcium concentration while preserving the integrity of the tip links.^[27]^ A holding current of +10 nA was continuously applied to counteract the diffusive release of the calcium chelator from the electrode, to which we added either a step or a succession of descending and ascending ramps of iontophoretic current. The steps had a magnitude of varying between −80 nA and−60 nA, while the ramps reached a minimal value between −90 nA and−50 nA. The current varied at the same rate on the descending and ascending ramps, within a range 5 − 8 nA ⋅ s^−1^, and the minimal current was maintained for 1 − 2 s before the ascending ramp started.

In a second series of experiments, we aimed at disrupting the tip links to evaluate their contribution to hair-bundle stiffness. In this case, the electrode was filled with EDTA at a concentration of 100 mM. We applied current steps of −100 nA on top of a resting current of +10 nA. In three cells, reducing the amplitude of the step to about −35 nA allowed to maintain the integrity of the tip links, adding data to those obtained with iontophoresis of triphosphate to evaluate the effects of reducing the calcium concentration on a force-displacement relation.

### Signal generation and acquisition

All signals were generated and acquired under the control of a computer running a user interface programmed with the LabVIEW software (version 2011, National Instruments). Command signals were produced by a 16-bit interface card (PCI-6733, National Instruments). A second 16-bit interface card (PCI-6250, National Instruments) conducted signal acquisition. Sampling rates for signal generation and acquisition varied within the range 2.5 − 25 kHz. All signals were conditioned with an eight-pole Bessel antialiasing filter (Model 3384, Krohn-Hite Corporation) at a lowpass cut-off frequency set at half the sampling frequency unless otherwise indicated.

### Analysis of spontaneous hair-bundle oscillations

Details of the analysis of the spontaneous oscillations have been published elsewhere.^[29]^ Briefly, the displacement, X(*t*), of an oscillating hair bundle was monitored over a duration of 10 − 30 *s*. The relation between its power spectrum, *C*(*f*), and frequency, *f*, was first smoothed by a moving average with a frequency window of 1 Hz and then fitted by the sum of two Lorentzians

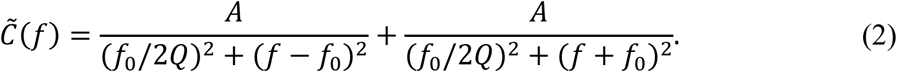

The fit provided values for the quality factor *Q*, the characteristic frequency *f*_0_, and parameter A, which is related to the root-mean-square magnitude 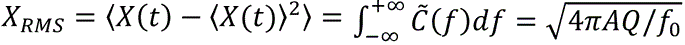 of the movement. In cases where it was bimodal, the probability density of hair bundle positions was fitted by the sum of two Gaussian functions. The fractional area under the Gaussian that was centered at a positive position with respect to the mean bundle position provided a functional estimate of the open probability 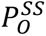 of transduction channels at steady state.

### Data analysis

Data were analyzed and visualized using MATLAB. Graphs were generated using MATLAB and Adobe Illustrator (Adobe).

### Statistical analysis

All numerical results are quoted as mean ± standard deviation (SD) over an ensemble of *n* independent measurements. Correlations were evaluated using Pearson’s correlation test, with a coefficient *r* that characterizes the sign and strength of correlation and a p-value that determines the statistical significance of the test. A two-tailed paired Student’s *t*-test was used to determine whether the difference between the means of two ensembles of paired measurements was statistically significant. G-power analysis ensured that sample sizes were sufficient to achieve at least 90% power at a 5% significance level. Statistical significance was defined as a p-value less than 0.05, with significance levels indicated as p < 0.05 (*), p < 0.01 (**), p < 0.001 (***). The results of statistical tests are listed in Table S2 (Supplementary Information).

## Funding

Fondation pour l’Audition grant FPA RD 2020-7 (PM)

Labex Cell(n)Scale grant ANR-11-LABX-0038 (PM)

Labex Cell(n)Scale grant ANR-10-IDEX-001-02 (PM)

Université Paris Sciences et Lettres and doctoral school “Physique en Île-de-France” grant (AJ)

## Competing interest

The authors declare that they have no conflict of interest.

## Author Contributions

P.M. supervised the study. P.M. and A.J. designed the experiments. A.J. and L.S. performed the experiments. A.J. and P.M. analyzed the data and wrote the manuscript.

## Data availability statement

All data needed to evaluate the conclusions in the paper are present in the paper and/or the Supplementary Information. Datasets for all the figures in the Main Text and Supplementary Information can be found on Zenodo (https://doi.org/10.5281/zenodo.14452874). Additional data related to this paper may be requested from the corresponding author.

Received: ((will be filled in by the editorial staff))

Revised: ((will be filled in by the editorial staff))

Published online: ((will be filled in by the editorial staff))

## Supporting Information

Supporting Information is available from the Wiley Online Library or from the author.

## Supplementary Table

**Table S1.**
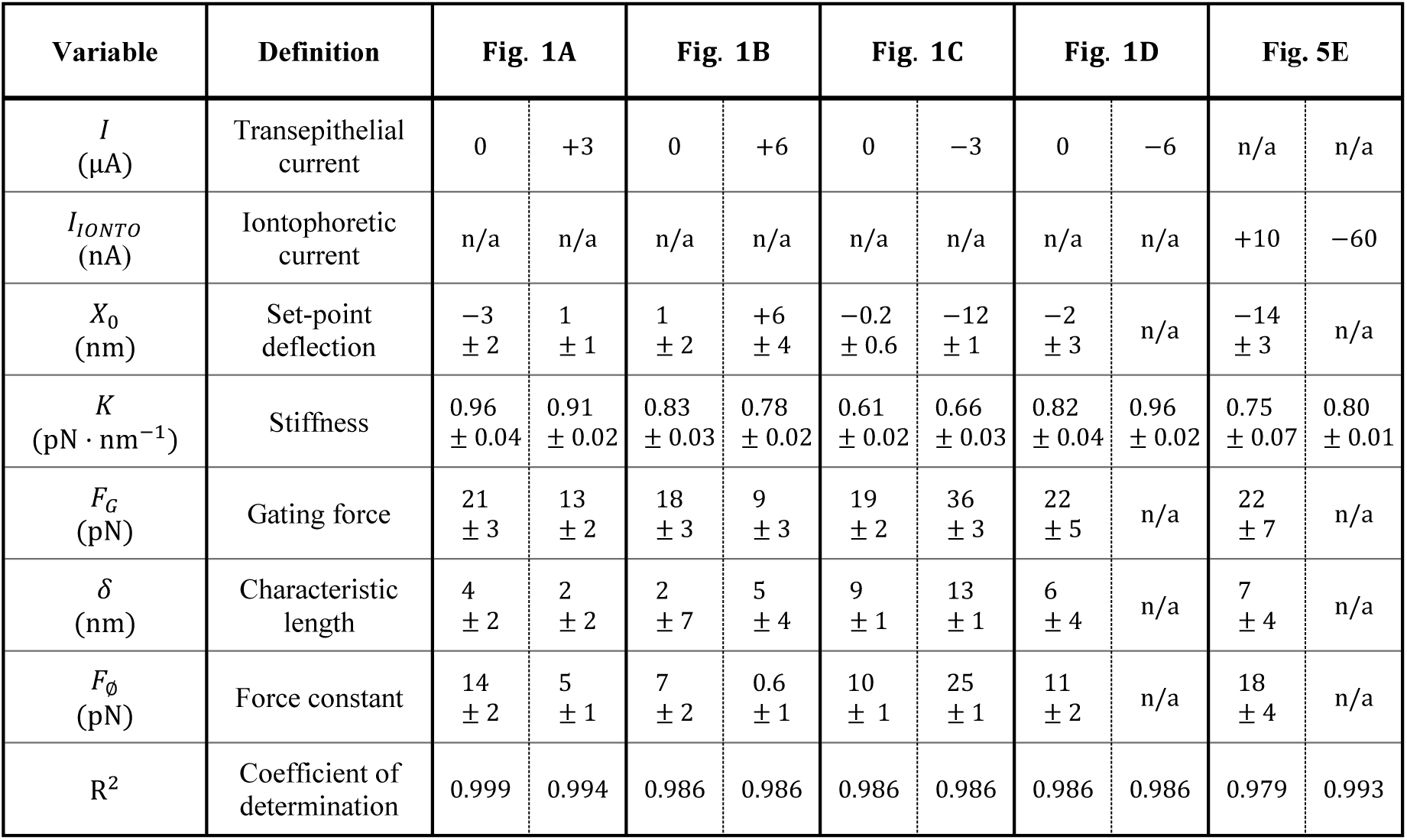
Parameter values with 95% intervals for fits of force-displacement relations with Equation 1.

**Table S2.** Statistical significance and power in two-tailed paired Student’s *t*-test.

|  | Fig. 5F |  | Fig. 6C |  | Fig. 7D |  |
| --- | --- | --- | --- | --- | --- | --- |
| Number of replicates | *n* = 8 cells in 6 frogs |  | *n* = 4 cells in 3 frogs |  | *n* = 8 cells in 6 frogs |  |
| Variable | Max. friction force, $\phi_{MAX}$ [pN] | | Rise time, $\tau_{10-90}$ [ms] | | Max. friction force, $\phi_{MAX}$ [pN] | |
| Test | Start (1) vs Pre-drop (2) | Start (1) vs Pre-drop (2) | Onset:$\phi_{MAX}$(1) vs $\bar{X}$ (2) | Offset:$\phi_{MAX}$(1) vs $\bar{X}$ (2) | Start (1) vs Pre-drop (2) | Start (1) vs Post-drop (2) |
| Mean (1) | 10.2 | 10.2 | 180 | 412 | 10.1 | 10.1 |
| SD (1) | 1.7 | 1.7 | 55 | 118 | 1.3 | 1.3 |
| Mean (2) | 13.2 | 12.1 | 487 | 100 | 12.1 | 8.2 |
| SD (2) | 2.2 | 1.6 | 64 | 44.1 | 1.6 | 1.19 |
| Effect size | 1.7 | 1.6 | 5.6 | −2.6 | 1.6 | −1.4 |
| Statistical power | 0.98 | 1.00 | 1.00 | 0.92 | 0.97 | 0.92 |
| p-value | $2 \cdot 10^{-2}$ | $1.5 \cdot 10^{-2}$ | $1.5 \cdot 10^{-2}$ | $1.5 \cdot 10^{-2}$ | $1.5 \cdot 10^{-2}$ | $5 \cdot 10^{-3}$ |

## Supplementary Figures

**Figure S1:**
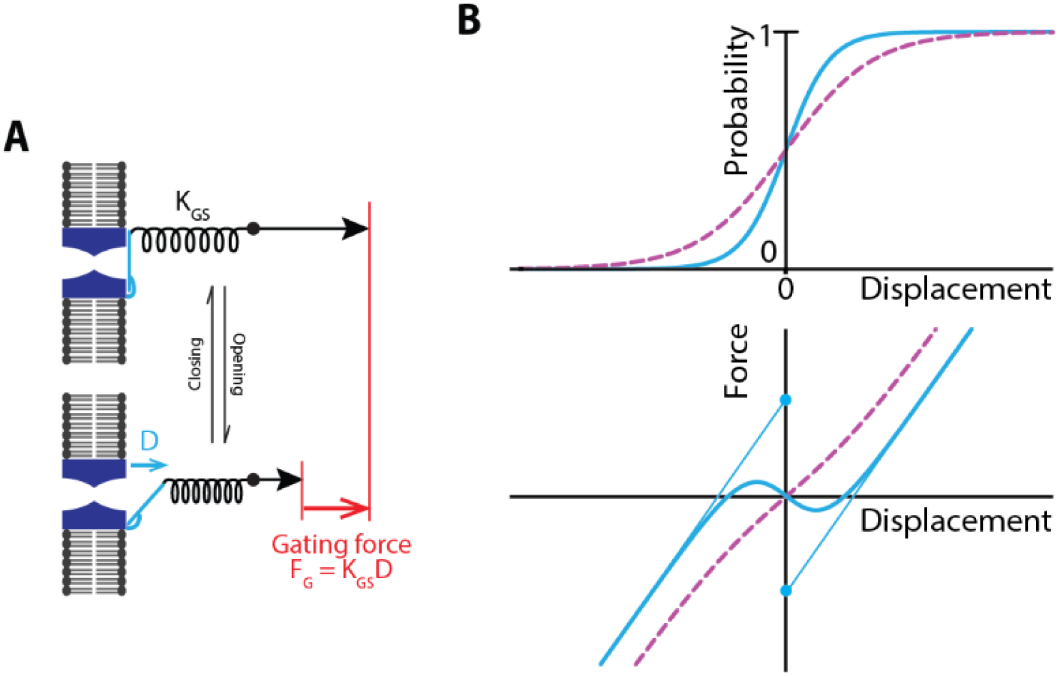
Gating-spring model of mechanoelectrical transduction. **(A)** The concept of gating force. When the transduction channels switch between their open and closed state, tension in the gating springs is reduced (black arrows); this mechanical correlate of channel gating is called the gating force, *F*_*G*_ = *K*_*GS*_ ⋅ *D* (red arrow), where *K*_*GS*_ is the combined stiffness of all the gating springs operating in parallel in the hair bundle and *D* is the gating swing of each channel. **(B)** Electrical and mechanical responses to hair-bundle displacement. Top: sigmoidal relationship of channel open probability to displacement in the case of weak (dashed magenta line) and strong (solid blue line) gating force. The larger the gating force, the steeper the sigmoid and the more sensitive the hair cell is to displacement of its hair bundle. Bottom: The force–displacement relationship displays a shallow nonlinearity when sensitivity is low (dashed magenta curve) and a strong nonlinearity when sensitivity is high (smooth blue curve). The gating force is given by the vertical shift between the two linear limbs of the force-displacement relation; it is here shown in the case of high sensitivity as the distance between the two disks.

**Figure S2:**
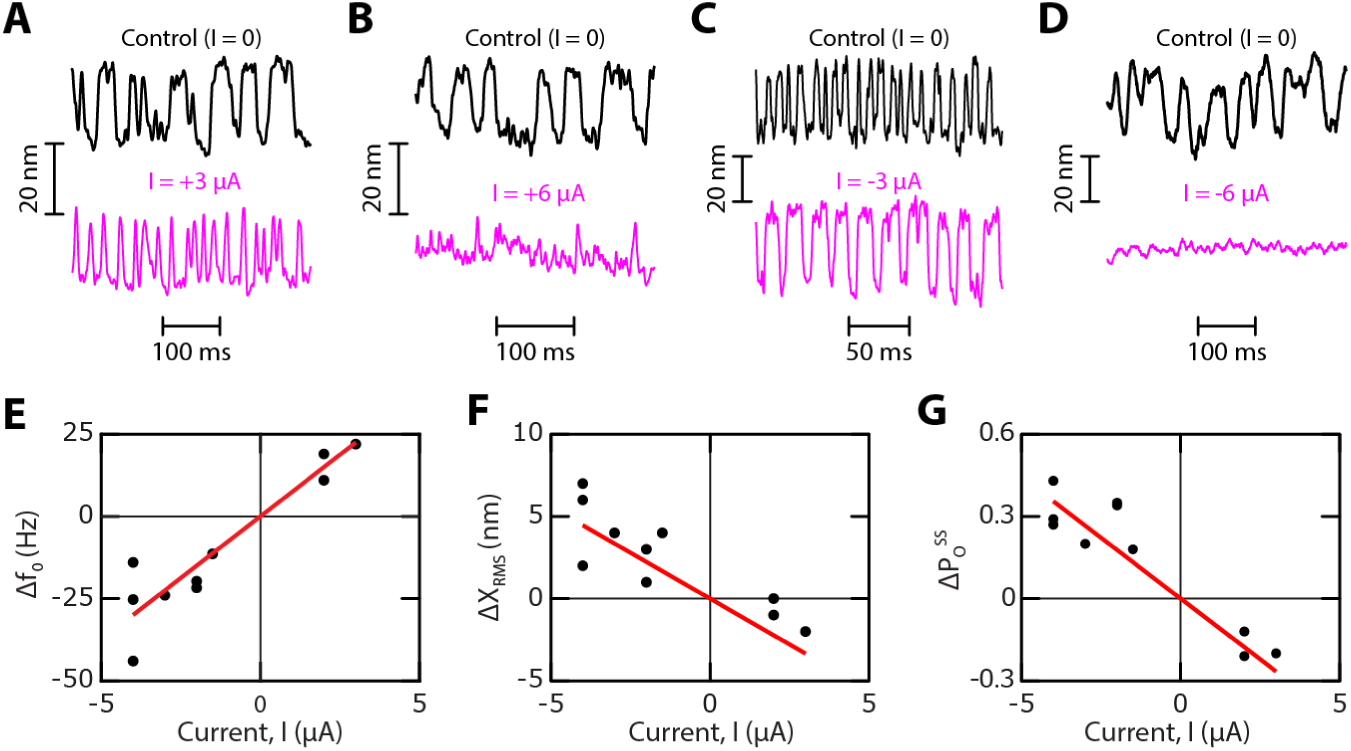
Effects of a transepithelial current on spontaneous hair-bundle oscillations. Under control conditions (*I* = 0), over an ensemble of *n* = 10 oscillatory hair bundles, the oscillations had a characteristic frequency *f*_0_ = 22 ± 14 Hz, a root-mean-squared (RMS) amplitude X_*RMS*_ = 13 ± 4 nm and their bimodal position histograms were associated with a steady-state open probability for the transduction channels 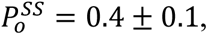 thus close to 1/2. These characteristics are remarkably similar to those reported earlier with hair cells from a different frog species.^[26,29]^ **(A-D)** Representative examples of spontaneous hair-bundle oscillations under control conditions (*I* = 0; top) and in the presence of a transepithelial current (bottom) *I* = +3 μA (A), *I* = +6 μA (B), *I* = −3 μA (C), and *I* = −6 μA (D). **(E)** The change, Δ*f*_0_, of the oscillation frequency displayed a strong positive correlation to the transepithelial current (Pearson correlation *r* = +0.9; p-value = 10^−4^; *n* = 10). **(F)** The change, *Δ*X_*RMS*_, of the RMS amplitude of spontaneous oscillations displayed a strong negative correlation to the transepithelial current (Pearson correlation *r* = −0.9; p-value = 1 · 10^−3^; *n* = 10). **(G)** The change, 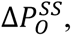 of the open probability at steady state displayed a strong negative correlation to the transepithelial current (Pearson correlation r = −0.9, p-value = 1 · 10^−4^, *n* = 10). In (A), the transepithelial current evoked an increase of the oscillation frequency from 17 Hz to 39 Hz, a decrease of the RMS amplitude of oscillation from 7 nm to 5 nm and a decrease of the steady-state open probability from 0.5 to 0.3. Conversely in (B), the frequency of oscillation decreased from 49 Hz to 25 Hz, the RMS amplitude of oscillation increased from 11 nm to 15 nm and the steady-state open probability increased from 0.4 to 0.6. In the presence of a large positive current (*I* = +6 µA; panel B), the hair bundle only produced irregular spiky movements in the positive direction: the transduction channels were closed most of the time and evinced brief openings. Instead, a large negative current (*I* = −6 µA; panel D) completely abolished spontaneous oscillations. Red lines in (E-G) correspond to proportional fits of respective slopes 7 Hz ⋅ μA^−1^(R^2^ = 0.86; panel E), −1.1 nm ⋅ μA^−1^ (R^2^ = 0.6; panel F), and −0.08 μA^−1^ (R^2^ = 0.83; panel G).

**Figure S3:**
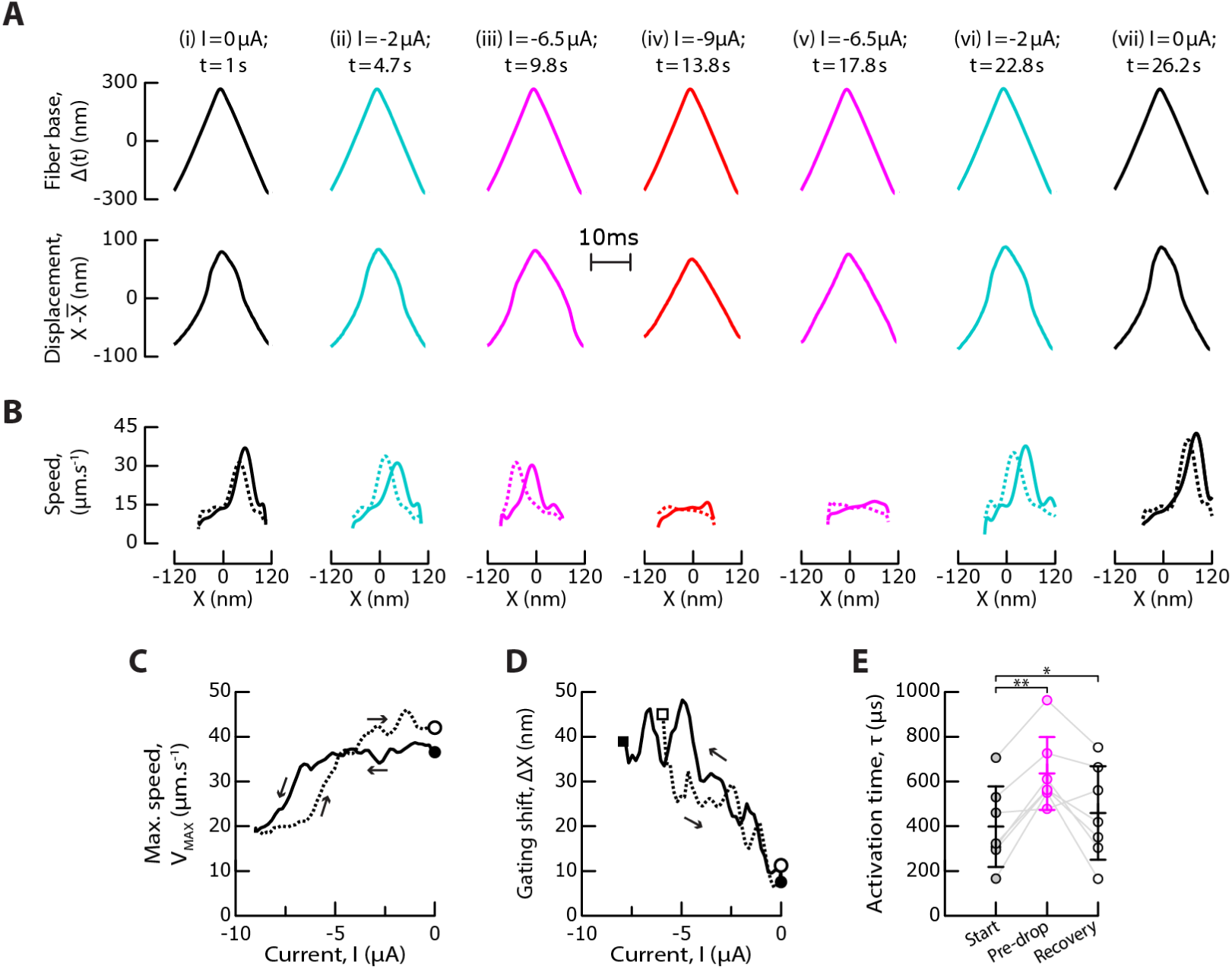
Effect of a transepithelial current on the channel-activation time. Same data as in Figure 5A-E, to which this figure is associated. The time course of successive ramps of descending and ascending transepithelial current can be found in Figure 5A. **(A)** Fiber-base displacement, *Δ* (*top*), and hair-bundle displacement, X (*bottom*), as a function of time for one stimulus cycle at time points and values of the transepithelial current indicated at the top of each time trace. **(B)** Hair-bundle speed, |*d*X/*dt*|, on the positive (*solid line*) and negative (*dashed line*) half cycles of mechanical stimulation as a function of displacement, X. The distance between the two peaks, *Δ*X, defines the gating shift. **(C-D)** Maximum speed, *V*_*MAX*_(C), and gating shift, *Δ*X (D), as a function of the transepithelial current, *I*, which varied in time from start (black disk) to finish (white disk). Arrows indicate the direction of circulation along the curve. Solid and dashed lines correspond to the descending and ascending ramps of transepithelial currents, respectively. In panel (D), a black square indicates the current at which the signatures of gating compliance and gating friction vanished on the descending ramp; bellow this current, the gating shift is no longer defined. Conversely, a white square indicates recovery of the signatures of gating compliance and gating friction on the ascending ramp. **(E)** For the same ensemble of hair bundles as in Figure 5, the distributions of the estimated activation time of the transduction channel, *τ* = ΔX/(2 *V*_*MAX*_), are shown at the start of the experiment (*I* = 0, black), before the drop of the maximal friction force (magenta, see Figure 5F), and at the end of the experiment (*I* = 0, white). Error bars: mean ± SD. Statistical significance was assessed using a paired t-test with, from p-values from left to right: 0.0014, 0.021 *p-value ≤ 0.05, **p-value ≤ 0.01.

## Legends for supplementary movies

**Movie S1 (separate file): Transition between states of strong and weak gating force.** Movie associated to Figure 3A-E. The force-displacement cycle in response to triangular mechanical stimulation of the hair bundle (top) is shown over time as the transepithelial current follows successive descending and ascending ramps (bottom). The friction force, *ϕ*(X), at each displacement X of the hair bundle is color coded as indicated on the right; it is given by the half-height of the force-displacement cycle. To obtain smooth videos, the force-displacement cycles were here averaged over 15 cycles, instead of 10 cycles in the data shown in Figure 3.

**Movie S2 (separate file): Kinetics of the transition between states of strong and weak gating force.** Movie associated to Figure 4A-B. The force-displacement cycle in response to triangular mechanical stimulation of the hair bundle (top) is shown over time before, during and after application of a negative step of transepithelial current (bottom). The friction force, *ϕ*(X), at each displacement X of the hair bundle is color coded as indicated on the right; it is given by the half-height of the force-displacement cycle. The force-displacement cycles were here averaged over 5 cycles.

**Movie S3 Transition between states of strong and weak gating force upon a decrease of extracellular calcium concentration.** Movie associated to Figure 5A-D. The force-displacement cycle in response to triangular mechanical stimulation of the hair bundle (top) is shown over time as an iontophoretic current trough a pipette containing a calcium chelator and positioned near a hair bundle follows successive descending and ascending ramps (bottom). The friction force, *ϕ*(X), at each displacement X of the hair bundle is color coded as indicated on the right; it is given by the half-height of the force-displacement cycle. To obtain smooth videos, the force-displacement cycles were here averaged over 15 cycles, instead of 10 cycles for the data shown in Figure 5.

## Notes

### Competing Interest Statement

The authors have declared no competing interest.

### Summary of Updates

- A new figure has been added to present a conceptual model accounting for the experimental findings; - Figures 3, 4, 5, and 7 have been tweeked to improve their readability, - The Discussion (now called 'Conclusion') has been expanded; - The Introduction has been revised.

